# Thermodynamic coupling between neighboring binding sites in homo-oligomeric ligand sensing proteins from mass resolved ligand dependent population distributions

**DOI:** 10.1101/2022.03.19.484990

**Authors:** Weicheng Li, Andrew S. Norris, Katie Lichtenthal, Skyler Kelly, Elihu C. Ihms, Paul Gollnick, Vicki H. Wysocki, Mark P. Foster

## Abstract

Homo-oligomeric ligand-activated proteins are ubiquitous in biology. The functions of such molecules are commonly regulated by allosteric coupling between ligand binding sites. Understanding the basis for this regulation requires both quantifying the free energy ΔG transduced between sites, and the structural basis by which it is transduced. We consider allostery in three variants of the model ring-shaped homo-oligomeric *<u>t</u>rp* <u>R</u>NA binding <u>a</u>ttenuation <u>p</u>rotein, TRAP. First, we developed nearest-neighbor statistical thermodynamic binding models comprising microscopic free energies for ligand binding to isolated sites ΔG_**N0**_, and for coupling between one or both adjacent sites, ΔG_N1_ and ΔG_N2_. Using the resulting partition function (PF) we explored the effects of these parameters on simulated population distributions for the 2^N^ possible liganded states. We then experimentally monitored ligand-dependent population shifts using conventional spectroscopic and calorimetric methods, and using native mass spectrometry (MS). By resolving species with differing numbers of bound ligands by their mass, native MS revealed striking differences in their ligand-dependent population shifts. Fitting the populations to a binding polynomial derived from the PF yielded coupling free energy terms corresponding to orders of magnitude differences in cooperativity. Uniquely, this approach predicts *which* of the possible 2^N^ liganded states are populated at different ligand concentrations, providing necessary insights into regulation. The combination of statistical thermodynamic modeling with native MS may provide the thermodynamic foundation for a meaningful understanding of the structure-thermodynamic linkage that drives cooperativity.

**TOC Figure (draft):** TOC Figure.
Ligand (Trp) binding to multiple sites on homo-oligomeric ring-shaped proteins like TRAP alters their functional states. Homotropic cooperativity is expected to alter the activation pathway in response to cellular ligand concentration. In the presence of positive nearest-neighbor cooperativity, ligand binding is favored at adjacent sites, whereas in the absence of cooperativity, a random “Normal” distribution is expected.

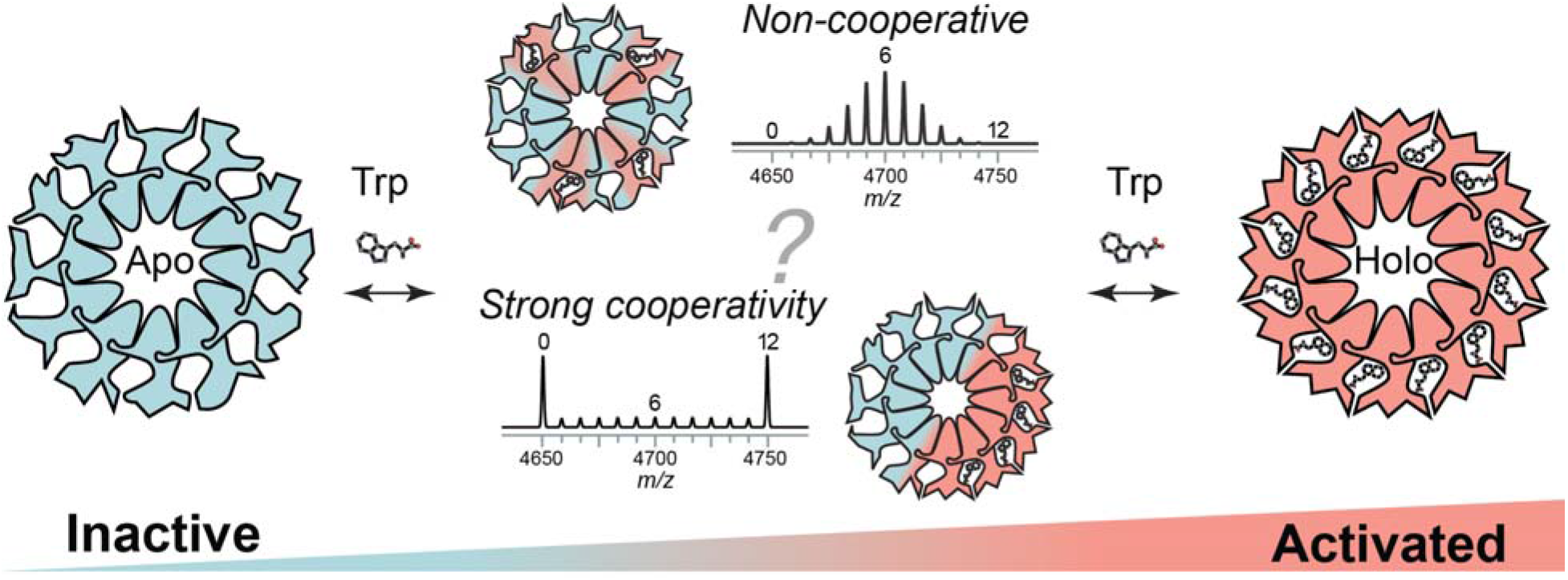

## Introduction

### Homotropic allostery is widespread in biology

Homotropic cooperativity describes a ubiquitous process in biological regulation. It arises commonly in proteins that form symmetric homo-oligomeric assemblies whose biological activity is modified by ligands.^1,2^ In such symmetric assemblies, even though the binding sites are identical, ligand binding affinities depend on the liganded state of the oligomeric protein; that is, the affinity *K* of a ligand for a receptor with no bound ligands will be different for a receptor that already has one or more bound ligands.^3^ This difference in affinity arises through “allosteric” interactions between the ligand binding sites, which alter the free energy change upon additional ligand binding.^4,5^ Such thermodynamic changes are most often associated with structural changes that accompany ligand binding, though changes in protein dynamics have also been implicated.^6,7^ A large proportion of biological processes are regulated by homooligomeric ligand binding proteins, from hemoglobin^8^ to p97 ATPase^9^. Therefore, to understand biological regulation we must first understand the linkage between ligand-induced structural changes and changes to protein-ligand free energy landscapes.^10^

### TRAP protein rings as models of homotropic allostery

We explore homotropic cooperativity in the homo-oligomeric *<u>t</u>rp* <u>R</u>NA binding <u>a</u>ttenuation <u>p</u>rotein TRAP, which plays a central role in regulating tryptophan (Trp) production in Bacilli.^11^ TRAP was originally identified in *Bacillus subtilis* (*Bsu*) as a 76-residue protein that assembles into homooligomeric complexes. Upon binding multiple Trp ligands, TRAP is activated to bind specific RNA sequences in the 5’ untranslated region of the *trp* operon mRNA.^12,13^ Crystal structures of TRAP from *Bsu*, *B. stearothermophilus* (*Bst*), *B. licheniformis*, and *B. halodurans* (*Bha*) revealed undecameric (11mer) and dodecameric (12mer) rings with binding sites for Trp in the interfaces between each of the protomers (Figure 1A-C). Trp-dependent sequence-specific RNA recognition is achieved via base contacts from amino acids located on loops that are flexible in the absence of Trp and become structured upon Trp binding; these loops thus mediate heterotropic Trp-RNA cooperativity.^13^ The same loops that mediate RNA binding also are points of contact between neighboring Trp binding sites, providing a structural scaffolding for mediating site-site interactions (Figure 1D).

**Figure 1.**
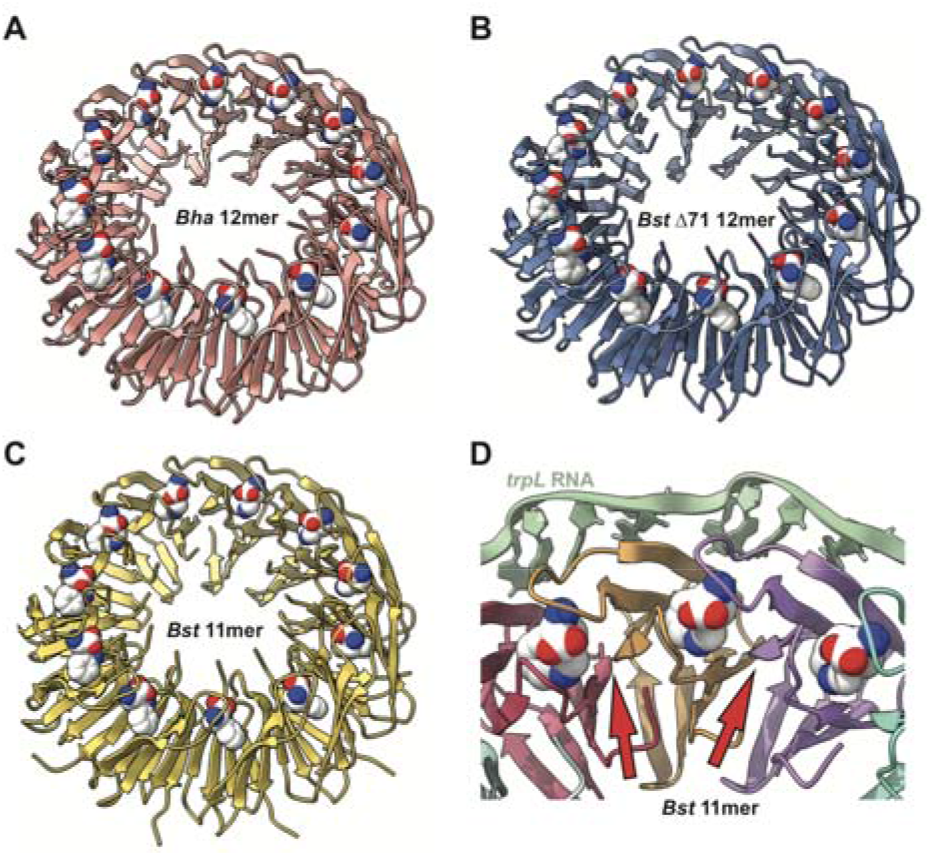
TRAP forms ring-shaped homo-oligomers to bind tryptophan and RNA. (A-C) Crystal structures of Trp-bound TRAP revealed 12mers for Bha TRAP (A; 3zzl), and for an engineered Bst TRAP-Δ71 (B; 3zzs), while wild-type Bst TRAP crystallized as 11mers (C; 1qaw). Bound Trp ligands are shown as spheres. (D) Crystal structure of Bst TRAP bound to an idealized trp leader RNA (green; 1C9S). Protein loops that become structured upon Trp binding participate in RNA recognition and mediate interactions between Trp binding sites (red arrows).

### Quantifying the detailed thermodynamics of ligand binding to homo-oligomeric proteins is difficult

Experimentally, binding affinities are usually quantified by measuring ligand-dependent changes in an observable that is proportional to the fraction of bound states *Y;* for example, a spectroscopic signal, or heat released upon binding. *Y* has the familiar form 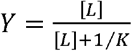, where [*L*] is the free ligand concentration and *K* is the equilibrium association constant. For a dodecameric protein with 12 binding sites, there are 2^12^ = 4096 possible configurations with 0-12 bound ligands. If the binding sites are identical and independent (i.e., in the absence of cooperativity), the populations of states with 0 to 12 bound ligands *L* follow a binomial distribution: 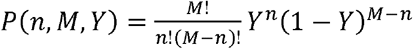, where *n* is the number of bound ligands, *M* is the total number of sites, and *Y* is the fraction of bound sites (i.e., the probability of sites being bound).^14,15^ Thus, for example, when enough ligand is present so that on average 1/12 of all sites are occupied (*Y* = 1/12 = 0.083), we find that states of the oligomer with one ligand bound are only slightly more abundant than those with none bound, or with two bound (0: 35%, 1: 38%, 2: 19%, 3: 6%) (Figure S1). If enough ligand is present to fill half of the sites (*Y* = 0.5), the most dominant states are those with six bound ligands, but other states are abundant according to a *normal* distribution, with coefficients 1:12:66:220:495:792:924:792:495:220:66:12:1 for 0-12 bound ligands. Thus, the resulting signal represents the average over all liganded states.

Moreover, the liganded states have configurational degeneracies: e.g., there are 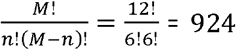 ways of binding 6 ligands to a symmetric homo-oligomer with 12 identical binding sites. The experimentally measured signal is thus the average over all liganded states *and* their configurations. Nevertheless, as long as the signal is proportional to *Y*, we can determine *K* by measuring how it changes with [L].

Cooperativity implies that *i* ligand binding events occur with *different* affinities *K_i_*. These differences result in altered population distributions, and how they respond to changes in ligand concentration. In the absence of cooperativity, populations can be expected to follow a purely statistical distribution that maximizes configurational entropy. At the extreme of infinitely positive cooperativity, under conditions of half maximal saturation (*Y* = 0.5), instead of a *normal* distribution we would observe a strict bimodal distribution: half of the oligomers will be fully bound, and half will be empty. Thus, cooperativity makes *some* configurations more favorable than others; *which* states are more favored depends on the *mechanism* and *magnitude* of the cooperative interactions. Thus, the *population distribution* encodes rich information about cooperative mechanisms that are convolved in the average metric *Y*.

A complicating factor in measuring population distributions is that cooperativity may distort the proportionality between the observable signal and fractional saturation *Y*. For example, in isothermal titration calorimetry (ITC) experiments, ligands may bind with differing affinities *K_i_*, but also with different enthalpies Δ*H*_i_; this makes complicates the use of heat as a metric for quantifying populations. Likewise, if a ligand bound at a site allosterically favors structural shifts in neighboring binding sites, subsequent ligand binding events as measured by circular dichroism (CD) might generate smaller structural changes and therefore lose proportionality to bound population.

### Native MS can measure population distributions

Native mass spectrometry (MS) has the potential to distinguish allosteric mechanisms by more directly probing bound-state distributions.^6,16–18^ Unlike aggregate measurements of average fractional saturation, native MS can directly probe the populations of states with different numbers of bound ligands.^18^ The mass (and mass-to-charge ratios, m/z) of a macromolecular complex is exactly determined by the molecular weight of its constituents. Thus, for an ionized macromolecule with *n* bound ligands, the signal will appear at the m/z dictated by the mass of the components and the charge on the ion. This direct proportionality between signal and ligand occupancy thus makes native MS well suited to characterize populations of states for molecules with multiple binding sites.

### Proper model selection is important

Finally, a severe confounding factor in understanding mechanisms of cooperativity is the selection of an appropriate model to extract thermodynamic parameters from binding data. In the absence of cooperativity, one may fit the signal change with the hyperbolic binding equation 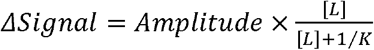 if the free ligand concentration is known or can be approximated, or the quadratic equation: 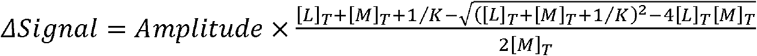 (*Equation* 1), if only total ligand and binding site concentrations [L]_T_ and [M]_T_ are known; *Amplitude* is the proportionality between fractional saturation *Y* and the signal change. However, these equations are inappropriate for cooperative binding since they only include one equilibrium constant *K*. Cooperativity is plainly evident if binding curves cannot be fit accurately with one of these equations.

Parsimonious approaches to data analysis demand adding only as many parameters as needed to produce a good fit to the binding data. A common approach is to use the Hill equation, which can generate sigmoidal binding curves but is uninformative about free energy differences between binding modes. Phenomenological multi-site independent binding models can also produce cooperative binding curves as well as different apparent affinities, yielding free energy differences between phenomenological binding modes.^19^

However, to decipher cooperativity mechanisms it is necessary to apply an appropriate physical binding model to the thermodynamic parameters. In the concerted Monod-Wyman-Changeux (MWC) model, all subunits of a homo-oligomer are posited to simultaneously exchange between distinct structures with different affinities for the ligand binding.^20^ In the sequential Koshland-Nemethy-Filmer (KNF) model, each subunit independently undertakes conformational changes upon ligand binding, altering the binding affinity for subsequent binding events.^21^ While each of these models are able to reproduce cooperative binding curves, they are unable to clarify which of the 2^N^ = 4096 possible liganded configurations of a 12mer are more favorable.

### Mechanistic insights from statistical thermodynamic models and native MS

Here, we explore the use of mechanistic nearest-neighbor (NN) lattice-based statistical thermodynamic models, along with native MS, to quantify and describe the cooperative behavior of three TRAP protein variants. We apply additive and non-additive NN models parametrized with free energies of binding to isolated sites, and microscopic free energies for communication between sites. We iterate over all possible 2^N^ liganded configurations of TRAP rings to develop statistical thermodynamic partition functions (PF) that describe the probabilities of each of the 2^N^ possible liganded states. We explore the effect of nearest-neighbor interaction energies on population distributions during simulated ligand titrations and examine the sensitivity of average site occupancy to the underlying population distributions.

We then measure Trp binding to TRAP variants using conventional bulk methods (CD, ITC), and using native MS. We find that native MS makes it possible to resolve and quantify states with 0 to *n* bound ligands, thereby allowing measurement of the statistical thermodynamic PF. Fitting of the populations of states with different numbers of bound ligands to a binding polynomial derived from the PF allows quantification of the microscopic free energy transduced between adjacent ligand binding sites. Armed with these tools, it is possible to predict *which* of the 2^N^ possible states are populated under different conditions, potentially distinguishing the *active* and *inactive* populations. Given the widespread occurrence in nature of homo-oligomeric protein assemblies,^1,2^ the approach outlined here for obtaining microscopic thermodynamic parameters should advance our understanding of structure-thermodynamic relationships in biological regulation.

## Materials and Methods

### Protein Preparation

Three TRAP variants were prepared for these studies: *Bha* TRAP, which forms predominantly 12mers, *Bst* TRAP, which forms predominantly 11mers, and *Bst* TRAP-Δ71, which forms predominantly 12mers.^22,23^ The proteins share the same protomer extinction coefficient (ε = 2980 M^-1^cm^-1^) at 280 nm as predicted using ProtParam (https://web.expasy.org/protparam/). Protein concentrations were determined using a Nanodrop 2000c spectrophotometer (Thermo Scientific) in droplet mode.

To produce *Bha* TRAP (WP_010897809.1), the coding region for the wild-type protein, in the pET17b plasmid (Novagen),^24^ was modified by the insertion at the C-terminus of sequences encoding the TEV protease cleavage site (ENLYFQ/G) followed by six histidine residues (a His-tag); after removal of the His-tag, the resulting protein consists of ENLYFQ on C-terminus (Figure S2B). Proteins were expressed in *Escherichia coli* BL21(DE3) cells and grown at 37°C in LB. At mid-log phase (OD_600_ = 0.4 to 0.6), 0.5 mM IPTG (isopropyl ß-D-1-thiogalactopyranosid) was added for induction of protein expression. Cells were harvested by centrifugation at 5,000 g for 5 min, resuspended in lysis buffer (500 mM NaCl, 20 mM NaPi, 100 mM imidazole, pH = 7.4, 1 mM benzamidine, 1 mM PMSF (phenylmethylsulfonyl fluoride)), and lysed in a French pressure cell at 10,000 psi. The lysate was centrifuged at 30,000 × g for 20 min and filtered (0.45 um cellulose acetate filters, Advantec) before loading onto 1mL HisTrap™ HP nickel column (GE Healthcare) on an AKTA fast protein liquid chromatography (FPLC) system. His-tagged *Bha* TRAP WT protein was eluted with an imidazole gradient from 100mM to 1M in elution buffer (0.1 to 1 M imidazole, 500 mM NaCl, 20 mM NaPi, pH = 7.4). The pooled protein sample was dialyzed into buffer A (100 mM NaCl, 50 mM NaPO_4_, pH 8.0) and treated with TEV protease at a mass ratio of 100:1 to cleave the C-terminal His-tag. The protease-treated fraction was then reloaded onto the 1mL HisTrap™ HP nickel column (GE Healthcare) and eluted with lysis buffer and the flow-through fraction collected. To remove bound Tryptophan (Trp), protein was denatured by dialysis for more than 6 hours at room temperature in a 3.5 kDa cut-off dialysis tubing membrane (Spectrum Labs) against 6 M guanidinium hydrochloride (Gdn-HCl) in buffer A. The fraction was then filtered using a 25mm minisart^®^ syringe filter (pore size: 0.2 μm, Sartorius) and further purified using reversed-phase high-performance liquid chromatography (RP-HPLC) on a 250 × 10 mm Proto 300 C4 column (Higgins Analytical) as previously described.^25^ The resulting protein fractions were lyophilized and stored at −80 °C until needed. The protein was refolded by dissolving the lyophilized pellet in 6 M Gdn-HCl buffer A at a concentration of ~1 mg ml^-1^, and gradually refolded by sequential dialysis against 3 M, 1.5 M, 0.75 M, and 0 M Gdn-HCl in buffer A in 3.5 kDa cut-off dialysis membrane (Spectrum Labs) for 12 hours per round at room temperature. The refolded protein was then purified by size-exclusion chromatography using a Hiload™ 16/600 Superdex™ 75 pg column (Sigma-Aldrich) in buffer A.

The gene of *Bst* TRAP WT (WP_033013997.1) and the C-terminal-five-residue-truncated *Bst* TRAP Δ71 were expressed from pET17b and pET9a vectors (Novagen, Inc.), respectively, as previously described.^16,22^ The proteins were separately expressed in *E. coli* BL21(DE3) cells, grown at 37°C in LB, and induced at OD_600_ = 0.4 to 0.6 with 0.5 mM IPTG. Cells were harvested by centrifugation, resuspended in lysis buffer B (100 mM K_2_HPO_4_, 50 mM KCl, 1 mM EDTA, pH = 8.0) plus 1 mM benzamidine and 1 mM PMSF (phenylmethylsulfonyl fluoride). The cells were broken in a French pressure cell at 10,000 psi. The lysate was clarified by centrifugation at 30,000 × g for 25 minutes. For every 5 mL of supernatant, 1 mL of 2% protamine sulfate was added. Then the lysate was stirred on ice for 30 minutes. The lysate was centrifuged at 30,000 × g for 25 minutes and the supernatant was heated to 80°C for 10 minutes followed by another centrifugation at 30,000 × g for 25 minutes. The supernatant was then dialyzed overnight against 50 mM Tris-HCl (pH = 8.0). Dialyzed proteins were loaded on Mono Q™ 10/100 GL column (GE Healthcare) in 50 mM Tris-HCl (pH 8.0). Proteins were eluted with a salt gradient from 0 M to 1 M NaCl in 50 mM Tris-HCl (pH 8.0). The same protocol, and buffers described above, were used to remove Trp and refold proteins to obtain apo (ligand-free) proteins, except that the 0.75M Gdn-HCl dialysis step was omitted during refolding, and that RP-HPLC purified *Bst* TRAP Δ71 was refolded at 55°C instead of at room temperature.

### Circular Dichroism (CD)

Tryptophan binding to TRAP was measured by CD spectroscopy at 25°C using a JASCO model J-715 spectropolarimeter with cell path length of 1 mm. Spectra were recorded between 226 nm and 230 nm using ~24 μM TRAP protomers (~2 μM ring) in 50 mM sodium phosphate buffer, pH 8.0, in the presence of increasing concentrations of Trp (0 to 300 μM). The change in CD signal at 228 nm, which shows the maximal difference between spectra of apo- and Trp-liganded TRAP, was used as a measure of tryptophan binding.^26^ The change in CD signal was fit with the quadratic single-site binding equation (*Eq*. 1).

### Isothermal Titration Calorimetry (ITC)

RP-HPLC-purified, refolded, and SEC-purified TRAP proteins were dialyzed extensively against buffer A. A stock solution of tryptophan was prepared by dissolving L-Trp powder (Sigma Aldrich, 93659) in the dialysate. Trp concentration was determined from its extinction coefficient at 280 nm of ε =5540 M^-1^ cm^-1^ using a Nanodrop 2000c spectrophotometer (Thermo Scientific) in droplet mode. The measured concentrations of Trp in the syringe and of TRAP protomers in the cell were 293 μM and 23 μM for *Bha* TRAP, 433 μM and 38 μM for *Bst* TRAP Δ71, and 645 μM and 74.7 μM for *Bst* TRAP WT, respectively. ITC thermograms were recorded on a MicroCal VP-ITC (Malvern Instruments) at 25 °C. A reference power of 30 μcal sec^-1^ and a stirring speed of 307 rpm were applied for all experiments. Deionized water was used in the reference cell. The software NITPIC was used to correct the baseline for each isotherm by iteratively optimizing the noise parameter *wrmsd* and then integrating the enthalpy at each titration point.^27^ The integrated heats showed bimodal binding behavior for *Bha* TRAP, and single mode for *Bst* Δ71 and *Bst* TRAP. The integrated enthalpy data were then fit with two-sites or one-site binding models using *itcsimlib.^5^* TRAP oligomer concentrations were corrected assuming a stoichiometry of 1 Trp per protomer based on the fitted stoichiometry n; corrected concentrations were 25.4 μM for *Bha* TRAP, 40.0 μM for *Bst* TRAP Δ71, and 48.6 μM for *Bst* TRAP WT. Confidence intervals in fitted parameters were estimated from 200 bootstrapped samples.

### Native Mass Spectrometry

Aliquots of purified proteins were dialyzed into 400 mM ammonium acetate (Sigma Aldrich, 431311). For *Bha* TRAP and *Bst* TRAP Δ71, the pH was adjusted to pH 8 by adding ammonium hydroxide (~1M NH_3_ in H_2_O, Sigma-Aldrich). Dialyzed protein concentrations (~ 60 μM protomers) were measured as above (ε = 2980 M^-1^cm^-1^) at 280 nm using a Nanodrop 2000c spectrophotometer (Thermo Scientific) in droplet mode. The Trp was dissolved in 400 mM ammonium acetate and the Trp concentration was measured as above (ε = 5540 M^-1^ cm^-1^) at 280 nm^28^. Samples for each titration point were prepared freshly in parallel by mixing the dialyzed protein solutions with serial dilutions from a 1 M Trp stock solution to the indicated ligand concentrations (0 to 140 μM).

Native mass spectrometry (MS) experiments were performed at room temperature (~25°C) on a Q Exactive Ultra-High Mass Range (UHMR) Orbitrap mass spectrometer (Thermo Fisher Scientific) modified to allow for surface induced dissociation, similar to that previously described.^29^ Emitters were pulled from borosilicate filament capillaries (OD 1.0 mm, ID 0.78 mm, Sutter Instrument) in-house on a P-97 Flaming/Brown Micropipette Puller (Sutter Instrument) that is fitted with a thermometer placed near the heating element. The temperature near the heating element was stabilized below 30 °C between each pull, and the same tip pulling parameters were used for each titration set. Each sample was loaded into an emitter using gel loading tips (Genesee Scientific, 0.5-10μl Ultra Micro Gel Tip) to avoid contamination, and then placed on a Nanospray Flex ion source (Thermo Fisher Scientific) equipped with a platinum wire to allow for static nano electrospray ionization. To obtain a stable total ion current, the spray voltage was initially ramped to about 1.4 kV and then gradually decreased to between 0.6 – 0.9 kV and held constant. For all measurements, the following instrument tune settings were kept constant: capillary temperature 250°C, Source DC Offset 21 V, S-lens RF level 200, detector optimization to low *m/z*, ion transfer target high *m/z*, injection flatapole DC 5 V, inter flatapole lens 4 V, bent flatapole DC 2 V, transfer multipole DC 0 V, C-trap entrance lens inject 1.8 V, HCD energy 1 V, HCD field gradient 200 V, and HCD cell pressure 4 (UHV Sensor ~3 - 4E-10 mbar). For most emitters, spectra were collected at resolutions of 6250, 12500 and 25000, as defined at 400 *m/z*. Each spectrum was obtained by averaging the same number of scans, and the injection time and averaged micro-scans were fixed. For *Bha* TRAP and *Bst* TRAP Δ71, each titration point was recorded three times using different emitters. For *Bst* TRAP, each titration point was recorded two times.

Data were manually examined using Xcalibur Qual Browser software (Thermo Fisher Scientific), and spectral deconvolution was carried out using UniDec.^30^ For quantification of Trp_n_-TRAP_m_ populations, all spectra from each titration set were initially deconvolved using MetaUniDec V4.4 and then later each spectrum was reprocessed using UniDec V5.0.1 to allow for the double deconvolution approach to be used to produce deconvolved zero-charge mass spectra where the proteoforms found in the apo TRAP spectrum from each titration are combined.^31,32^ The titration spectra collected at a resolution of 12,500 were used for the final deconvolutions. Custom deconvolution parameters were used for each protein titration set. For all sets, UniDec parameters were adjusted to a narrow *m/z* range around the dominant oligomer charge state distribution, the charge range was set to less than 11 charges surrounding the most abundant charge state; mass sampling was set every 1 Da, FWHM (full width at half maximum) was set to 0.8 Th, peak shape function was set to split Gaussian/Lorentzian; artifact suppression, point smooth width and mass smooth width were set to 0; charge smooth width was set to 1.5, native charge offset range was set to +/− 6, and *m/z* to Mass transform was set to interpolate. A Mass List Window width of +/− 1200 Da was used. A mass list for each protein was generated based on the expected protein mass, expected oligomeric state(s), and 0 to 13 tryptophan bound (including bound 13mer). For *Bst* TRAP and *Bst* TRAP Δ71 this list consisted of just the expected 11mer and 12mer oligomeric states, respectively, with variable Trp. For *Bha* TRAP, we found that in addition to the expected 12mer, overlapping 13mer charge states were present, so the mass list was set to include the 13mer with variable Trp. The respective deconvolved apo TRAP zero-charge mass spectrum was used as the double deconvolution kernel file. The areas for each species in the deconvolved spectra were extracted, sum normalized, and used for the model fitting. The area extraction window was set to +/− two times the standard deviation. The standard deviation was determined based on the average FWHM of the deconvolved apo and holo peaks using the simplified equation where FWHM divided by 2.35 equals the standard deviation.

Accurate determination of protein concentration is complicated for TRAP proteins because they lack encoded Trp residues and have low extinction coefficients as noted above. Before thermodynamic analysis of the MS titration data, protein concentrations were corrected by fitting the bound fraction at each Trp concentration with the quadratic independent sites model; [*L*]_t_ was assumed to be accurate, while [*M*]_t_ and the apparent dissociation constant *K*_d_ were fit parameters.^16^ This fit yielded the binding site concentration [*M*]_t_ that was in general about 33% higher than initial estimated concentration (Table S3, Figure S8).

### Fitting of population distributions with NN models

Following TRAP concentration correction, experimental data obtained from native MS were fit using an additive nearest-neighbor (NN-add) model for 12 binding sites via *itcsimlib^5^*, as described previously.^16^ Briefly, the relative abundance of Trp-TRAP ligand-bound species (13 in total) were determined from the integrated peak areas, as mentioned above. To fit this population distribution the NN-add model was parametrized with values for ΔG_0_ (intrinsic binding free energy) and ΔG_N2_ (binding free energy with two occupied neighbors, equal to 2ΔG_N1_). A binding partition function that includes all possible Trp_n_-TRAP_12_ configurations (2^12^ = 4096) was generated using *itcsimlib*. The statistical thermodynamic probabilities of each configuration were then computed at each experimental ligand and binding site concentration. The ST probability of each of 4096 configurations were summed and converted into the relative abundance of 13 ligand-bound species, permitting generation of a virtual titration dataset. Then the discrepancy between native MS experimental titration data and the virtual titration data was globally minimized by iteratively varying the parameters of ΔG_0_ and ΔG_N2_ using a Powell^33^ gradient-descent optimization algorithm. Uncertainties in fit parameters were obtained from the standard deviation of three independent experimental repeats. Each dataset was also fit using a nonadditive nearest-neighbor model (NN-nonadd) with three parameters, ΔG_0_, ΔG_N1_, and ΔG_N2_. Inclusion of the extra parameter did not significantly improve the quality of the fits, so data were analysed using the simpler NN-add model.

### Population simulations with NN models

Simulated population distributions for 12-mer proteins were computed using additive or nonadditive nearest neighbor (NN) statistical thermodynamic models, with the same ΔG_N2_ = −7.5 kcal mol^-1^ (binding free energy with two occupied neighbors) but varied NN cooperativity factor α (and β in NN-nonadd model), and therefore different ΔG_0_ (intrinsic binding free energy).^5^ In the NN-add model, given ΔG_0_ and α values, the statistical thermodynamic probability of each of Trp_n_-TRAP_12_ configuration (2^12^ = 4096), along with the probabilities of the four basic NN energy configurations (0, N0, N1, N2) were computed over a range of 280 ligand concentrations using *itcsimlib.^5^* From these trajectories we computed fractional saturation curves by summing the three ligand-bound states, N0, N1, and N2, as a function of free ligand concentration (Figure 2C; Figure S3). The relative abundance of species with 0 to 12 bound Trp was used to simulate gaussian peak shapes for each state (Figure 2B). A similar set of simulations was performed for NN-nonadd model, including all three thermodynamic parameters of ΔG_0_, ΔG_1_, and ΔG_2_ (Figure 3; Figure S5).

**Figure 2.**
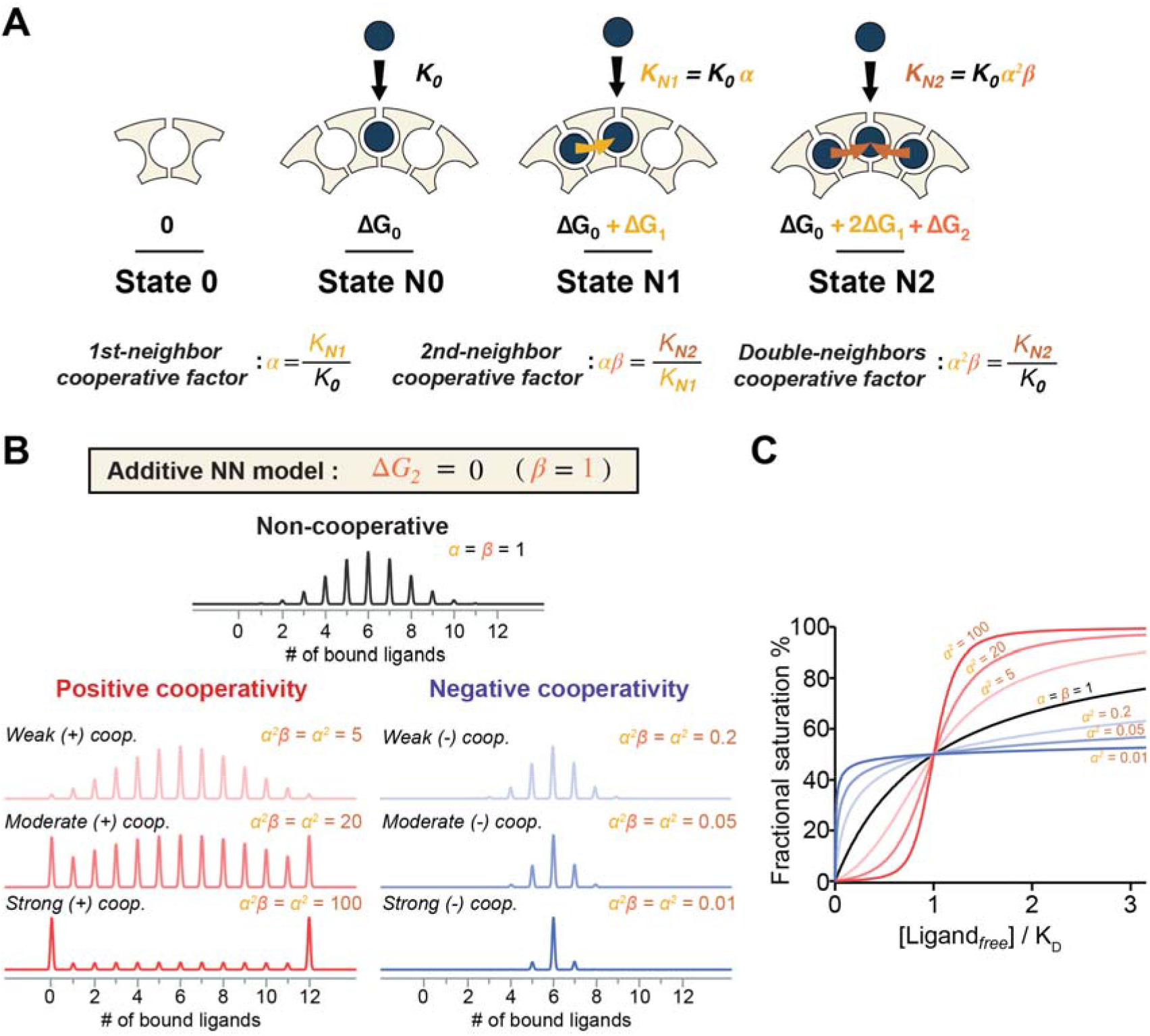
Nearest-neighbor models can explain and predict population distributions for cooperative ligand binding to ring-shaped oligomeric proteins. (A) In a nearest-neighbor (NN) model of cooperativity, four basic ligand binding states can be described by three energy parameters: an intrinsic free energy change for binding to sites with no occupied neighbors (ΔG_0_), a coupling free energy term contributed solely from each adjacent bound ligand (ΔG_1_), and a coupling free energy term contributed solely from the additive degree of two bound ligands (ΔG2). An empty site is defined as reference state (State 0) with ΔG_0_ = 0. Cooperativity from one bound neighbor (α) then can be defined by comparing the intrinsic equilibrium association constant (K_0_) to that with one bound neighbor (K_N1_) Cooperativity from having two bound neighbors (αβ) can be defined by comparing the two-neighbor binding (K_N2_) to one-neighbor binding (K_N1_). α and β values > 1 indicate stronger binding to sites with occupied neighbors. If K_N1_/ K_N0_ = K_N2_/K_N1_, i.e., ΔG_2_ = 0, β = 1, the coupling energy for two bound neighbors is the sum of each individual NN interaction, and the nearest neighbor effect is additive (NN-add); otherwise, it is non-additive (NN-nonadd). Cooperativity for binding with two neighbors (*α^2^β*) is defined by comparing the two-neighbor binding (K_N2_) to intrinsic binding (K_0_). (B) Effect of nearest-neighbor cooperativity on simulated population distributions for dodecameric rings with 0 to 12 bound ligands at half-saturation. Simulated populations for a non-cooperative binding (*α* = 1, black), equivalent to non-interacting sites, yields the expected binomial distribution. Population distributions arising from both positive (*α* > 1, red) and negative (< 1, blue) cooperativity were simulated using an additive NN model. In the presence of stronger positive cooperativity, as expected, the apo and holo populations dominate the distribution. In contrast, negative cooperativity favors states with half of the sites occupied. (C) Effects of NN cooperativity on fractional saturation of ligand binding sites. Positive cooperativity (red) generates sigmoidal binding curves whereas negative cooperativity (blue) generates steep but shallow curves; the non-cooperative binding curve is hyperbolic (black).

**Figure 3.**
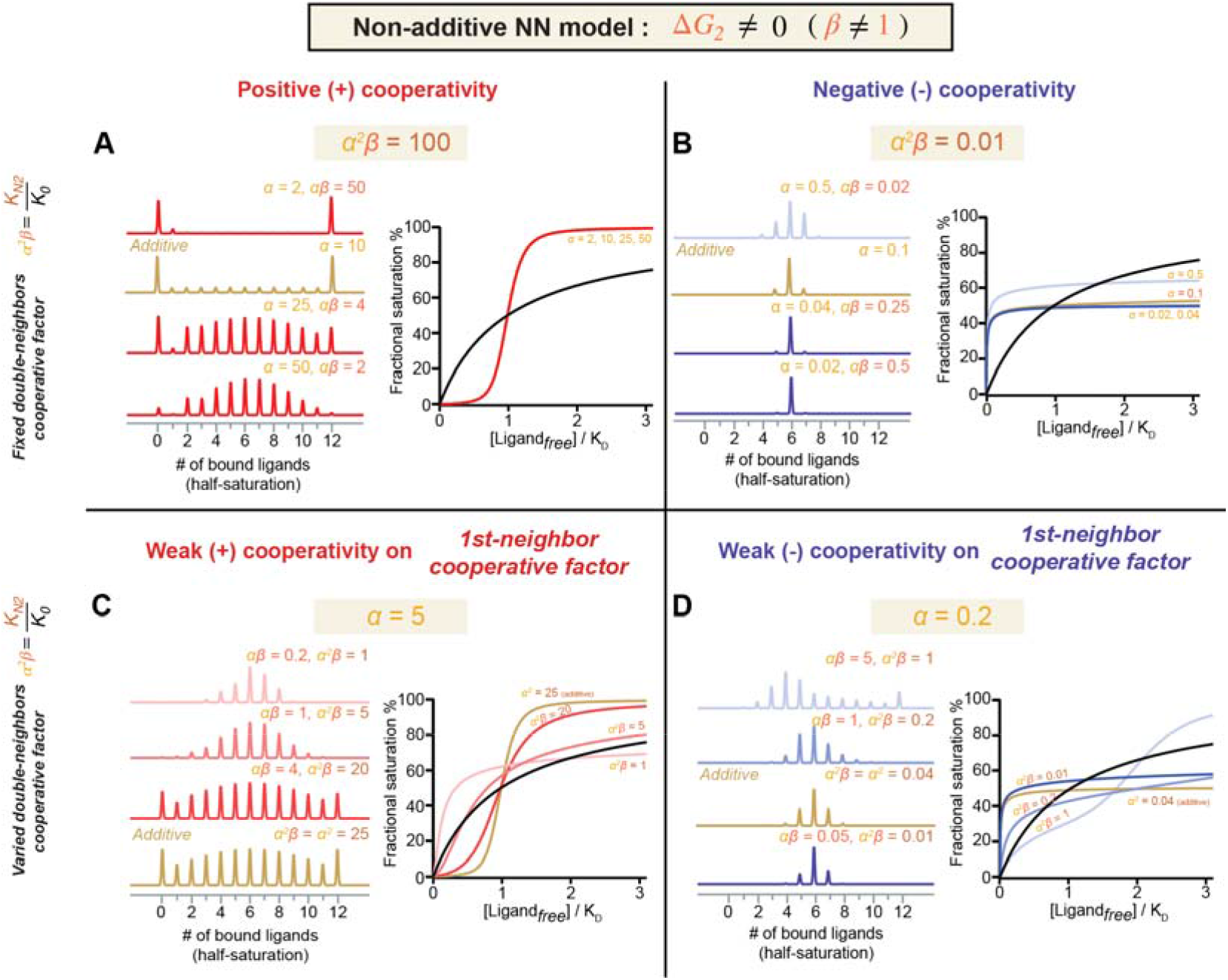
Simulated population for a cyclic-12mer bound to 0-12 ligands at concentrations that yield half maximal saturation, predicted by a non-additive nearest-neighbor (NN-nonadd) model with fixed or varied cooperativity strength, ^2^β. Fractional saturations and population distributions are shown for (A) positive cooperativity with fixed α^2^β = 100 and varied α, β; (B) negative cooperativity with fixed α^2^β = 0.01 and varied α, β; (C) positive cooperativity with fixed α = 5 and varied β; (D) negative cooperativity with fixed α = 0.2 and varied β.

## Results

### Nearest-neighbor statistical thermodynamic models can quantify cooperativity and provide mechanistic insights

Nearest-neighbor (NN) models posit that the thermodynamics of ligand binding to a site are most strongly influenced by the liganded states of the nearest-neighbor binding sites.^5^ Thus, the thermodynamics of binding to a given site can be parametrized using three microscopic free energy terms (ΔG_0_, ΔG_1_, ΔG_2_, Figure 2A), corresponding to free energies for binding to sites with no bound neighbors, and thermodynamic contributions from having one or two occupied neighbors, respectively; ΔG_1_ = ΔG_2_ = 0 in the absence of cooperativity. Therefore, four fundamental ligand binding states are: a) state-0, empty sites; b) state-N0 (ΔG_N0_ = ΔG_0_), bound sites with no occupied neighbor; c) state-N1 (ΔG_N1_ = ΔG_0_ + ΔG_1_), bound sites with one occupied neighbor; and d) state-N2 (ΔG_N2_ = ΔG_0_ + 2ΔG_1_ + ΔG_2_), bound sites with two occupied neighbors. The corresponding ligand binding affinity for each state is *k*_0_ = 1, 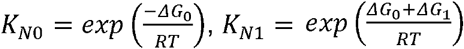, and 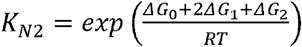, respectively. If the nearest-neighbor interactions are *additive*, i.e., ΔG_2_ = 0, the models are reduced to two free energy terms, with 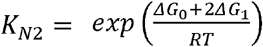. In addition, it is convenient to quantify “cooperativity” by comparing the affinities for ligand binding in various modes, such that α = K_N1_/K_N0_ > 1 quantifies the fold *increase* in affinity afforded by having the first occupied neighbor, while α < 1 quantifies the fold *decrease* in affinity from negative cooperativity. Likewise, we define αβ = K_N2_/K_N1_ to describe the *additional* fold change in affinity contributed from the second bound neighbor. Thus, α^2^β = K_N2_/K_N0_ quantifies the accumulated changes in affinity from both two occupied neighbors. With these parameters, NN models enable elaboration of a statistical partition function that enumerates the free energies and Boltzmann probabilities of all possible 2^N^ states by straightforward iteration and examination of their nearest-neighbor site occupancy.^5,34^

Application of an additive NN (NN-add) model to a cyclic homo-dodecamer results in a partition function (PF) 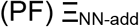 with 38 unique energetic terms, out of 2^12^ = 4096 configurations (*Equation 2*):

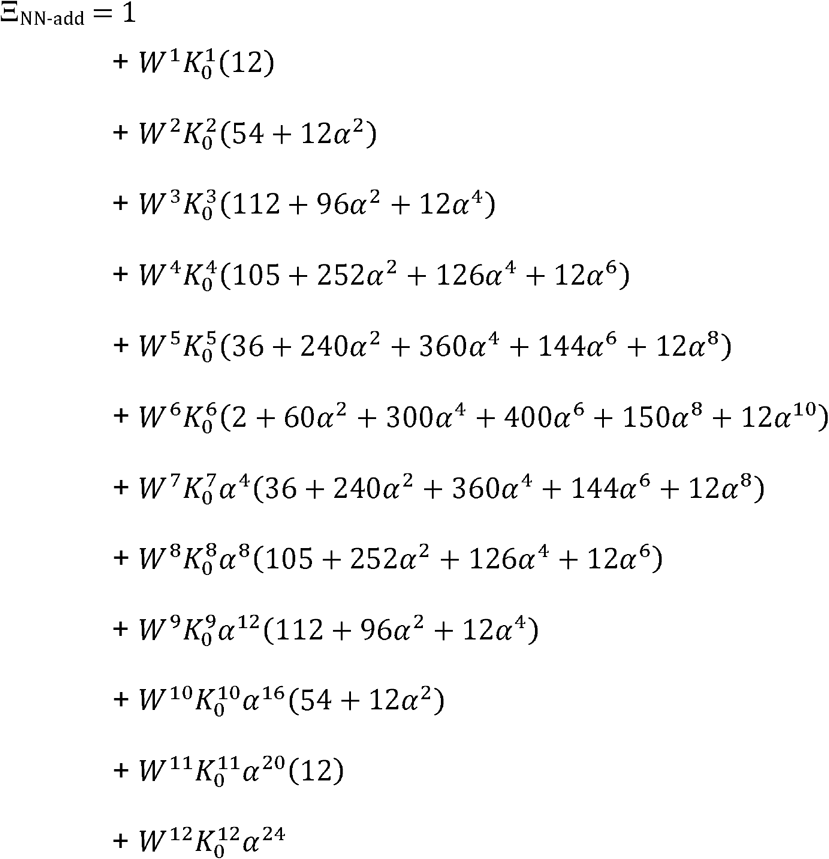

Here, *W* is the free Trp concentration and *K*_0_ and α are as defined above; generation of the PF was automated using *itcsimlib.^5^* Each row in Equation 2 corresponds to states with different numbers of bound ligands, from 0 to 12, with 1 assigned as the statistical weight of the state with zero bound ligands. For each term, the coefficients reflect the degeneracy of the energetic states. For example, there are 12 equivalent ways to configure 1 or 11 ligands on a 12-mer. For two bound ligands, there are 12 ways to arrange them next to each other, and modified by a cooperativity term α^2^, and 54 ways to arrange them in isolation, with no α^2^ term (Figure S3A). The exponent on the α terms reflects the number of nearest-neighbor interactions in that configuration, such that, for example, the half-bound state with the 400α^6^ term corresponds to configurations that arrange the six bound ligands with three pairs of interactions (e.g., Figure S3B).

With the NN-add partition constructed, it is possible predict the effect of cooperativity on the populations of each of the 4096 states in a ring-shaped homo-dodecameric proteins. As noted above, in the absence of cooperativity, at ligand concentrations that result in half of the sites being bound, the populations follow a binomial distribution (Figure 2B, black). Positive cooperativity increases the population of states with more bound ligands because they feature more nearest-neighbor interactions; this also results in more states with fewer sites bound, due to mass balance (Figure 2B, red graphs; Figure S4A). In the case of strong positive cooperativity, apo and holo states dominate the population distribution, with intermediate states being lowly populated. In contrast, negative cooperativity has the opposite effect, favoring half-bound configurations that spread out ligands over all available sites while minimizing NN interactions (Figure 2B, blue graphs; Figure S4B).

Nearest neighbor interactions may also be *non-additive*, meaning that the thermodynamic contributions from having two bound neighbors is not simply twice the contribution of having one bound neighbor (β ≠ 1). In this case, a non-additive nearest-neighbor (NN-nonadd) model is described by three independent energetic terms: ΔG_0_, ΔG_1_, and ΔG_2_ (Figure 2A). This formulation results in a partition function with 61 unique energy terms (Figure S5). The population distribution of states with 0-12 bound ligands were also simulated for a NN-nonadd model under conditions of half-maximal saturation (Figure 3) with a range of cooperativity strengths α and β. Compared to the additive model, we see that divergence from additivity (larger differences in ΔG_N1_ and ΔG_N2_) results in *asymmetry* in the population distributions. For example, more favorable ΔG_N2_ than ΔG_N1_ (or β > 1) favors more state-N2, depleting states with only a few ligands bound (Figure 3A, Figure S6A). *Detection of such asymmetry in the underlying distributions would be important for distinguishing mechanisms of allosteric communication*.

In addition to providing details of the underlying populations, NN models can be used to compute bulk properties for comparison with experimental measurements (Figure 2C). A plot of fractional saturation versus free ligand concentration generates the expected hyperbolic curve in the absence of cooperativity (black). Positive cooperativity produces the expected sigmoidal curves (red) and negative cooperativity yields compressed saturation curves (blue). Notably, different degrees of non-additivity may not be evident from bulk properties: for example, in the case of net favorable cooperativity with α^2^β = 100, though different values of α and β yield different underlying distributions, the saturation trajectories overlap (Figure 3A). Other combinations of parameters yield somewhat distinguishable saturation curves, but the underlying population distributions are much more informative of the underlying thermodynamics (Figure 3). Because ligand binding may alter the structure and function of oligomeric proteins (i.e., activating or repressing its function), such an analysis provides a thermodynamic link between the function of the protein and specific ligand-protein states.

### Cooperative Trp binding to multimers can be difficult to characterize using traditional bulk binding assays

To quantify Trp binding to oligomeric TRAP rings, we first used circular dichroism (CD) spectropolarimetry to measure its ellipticity at 228 nm, which reports on Trp binding (Figure 4A).^26^ CD measurements were conducted at 25°C with ~24 μM TRAP (2 μM rings) over a range of Trp concentrations. Binding curves were fit with the binding quadratic (*Eq*. 1), yielding apparent dissociation constants (*K*_d,app_). The three TRAP variants yielded similar apparent Trp binding affinities of 3-14 μM (Table S1), consistent with previous findings.^22^ Under these conditions, cooperative binding of Trp to the TRAP variants is not evident.

**Figure 4.**
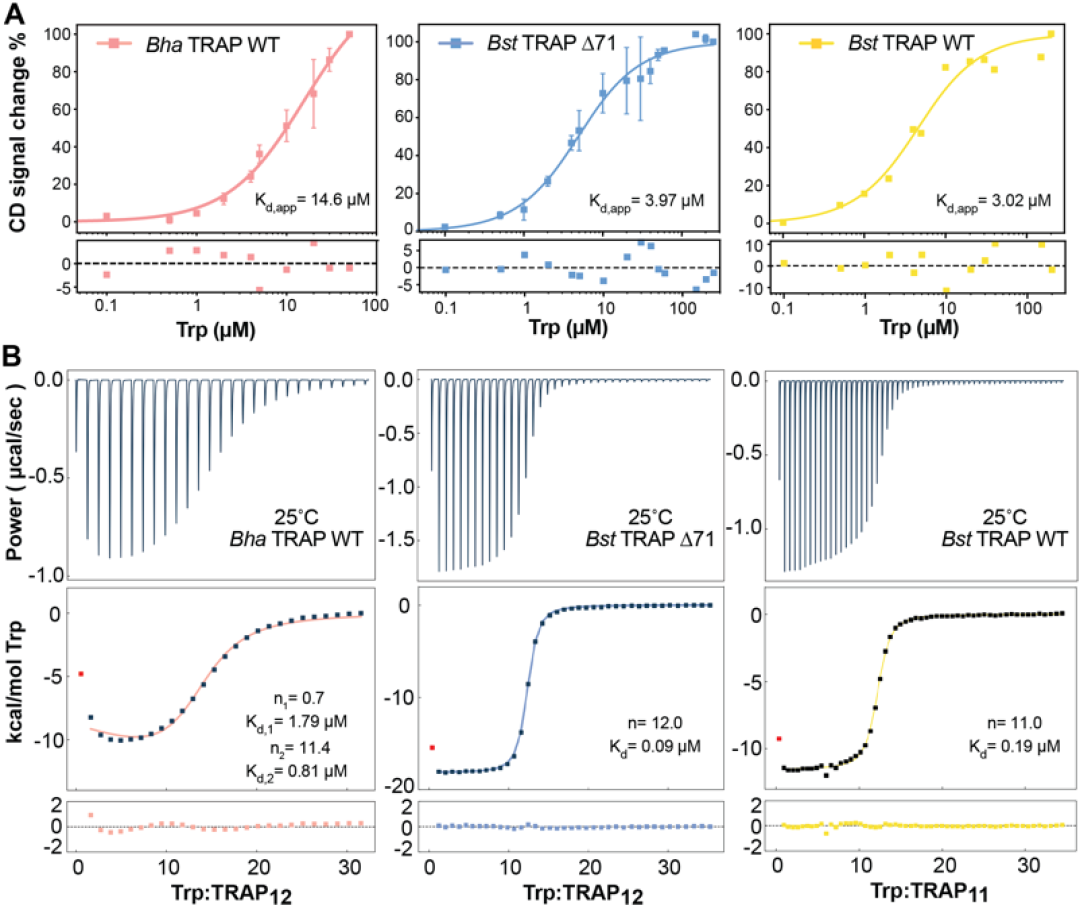
Bulk binding and phenomenological modeling provide limited information on cooperative ligand-binding to dodecameric Bha TRAP and Bst TRAP Δ71 (left, middle), or undecameric wild-type Bst TRAP (right). (A) The circular dichroism signal at 228 nm in 50 mM sodium phosphate buffer (pH = 8) was measured as a function of added Trp, and the data fit with the quadratic binding equation (Eq. 1; Table S1). CD data were acquired by using 2 μM TRAP with increasing Trp concentration from 0 to 300 μM. The fitted parameters of CD suggest all three proteins exhibit similar μM binding affinity. Homotropic cooperativity is not evident under these conditions. (B) ITC experiments titrating Trp into TRAP at 25°C in buffer A (see Methods) were fit with quadratic independent site models (Table S2). Bst TRAP Δ71 (middle) and Bst TRAP (right) can be well fitted with one-site binding model, in which 12mer Bst TRAP Δ71 (K_d_ = 0.09 μM) has two-times higher binding affinity than 11mer Bst TRAP (K_d_ = 0.19 μM). However, the data for Bha TRAP (left) is better fit to a two-sites binding model with improbable parameters in which the first Trp binds with K_d_ of 1.79 μM while the remaining eleven bind with a K_d_ of 0.81 μM.

Homotropic cooperativity can alter thermodynamic parameters other than binding free energy (i.e., *K*_d_), so we also studied Trp binding to TRAP using isothermal titration calorimetry (ITC), which directly measures enthalpy changes resulting from ligand binding. The binding thermograms for *Bst* TRAP12 Δ71 and *Bst* TRAP_11_ could be well fit with single-site (noncooperative) model, with comparable affinities (Figure 4B; Table S2). However, the binding thermogram for *Bha* TRAP_12_ is biphasic (Figure 4B), indicative of thermodynamically inequivalent binding events. Fitting of the thermograms for *Bha* TRAP to a two-site binding model yielded parameters with an initial (n ≈ 1) ligand binding event of *K*_d,1_ = 1.79 ± 0.30 μM, and subsequent 11 binding events with *K*_d,2_ = 0.8 ± 0.11 μM. Such parameters would suggest a binding model in which initial weaker binding of the first ligand fundamentally alters the structures of all other binding sites to favor binding, or alternatively, that all subsequent binding events occur in a defined sequential manner. These models describe drastically different modes of cooperativity but are indistinguishable by this method, which provides little insight into the mechanism by which thermodynamic communication occurs between sites. The results are also seemingly at odds with the CD experiments, which indicated weak cooperativity for *Bst* and *Bst* Δ71, but no cooperativity for *Bha* TRAP.

### Native MS enables quantification of TRAP states with different numbers of bound ligands

We used native mass spectrometry to quantity populations of Trp_n_-TRAP_m_ states over a range of Trp concentrations for *Bha* TRAP_12_, *Bst* TRAP_12_ Δ71 and *Bst* TRAP_11_ (Figure 5A). For each protein, titration samples were prepared in parallel by incubating the proteins in 400 mM ammonium acetate solutions with differing concentrations of Trp. For each oligomer, we obtained spectra with narrow charge distributions indicative of native structure.^35^ The most abundant charge states for each protein were 24+, 22+ and 21+ for *Bha, Bst*-Δ71 and *Bst* TRAP, respectively. Recorded spectra provided baseline resolution of signals with different numbers of bound Trp. Double deconvolution of the *m/z* spectra to include proteoforms ^32^ resulted in very clean zero-charge mass spectra, and reduced chi-squared values when fitting (Figure 5b; Figure S7). Spectra recorded in the absence of Trp contained signals corresponding to the apo oligomeric protein rings, and spectra at the highest Trp concentrations were dominated by the fully bound holo states, while intermediate states could be discerned at all non-zero Trp concentrations. Thus, these spectra enable us to resolve and quantify populations of states corresponding to the ligand-free apo states, the fully liganded holo states, and intermediate species with differing numbers of bound Trp ligands. These populations can be directly compared to the populations predicted by NN models (Figures 2, 3), and used to determine mechanisms and thermodynamic cooperativity parameters.

**Figure 5.**
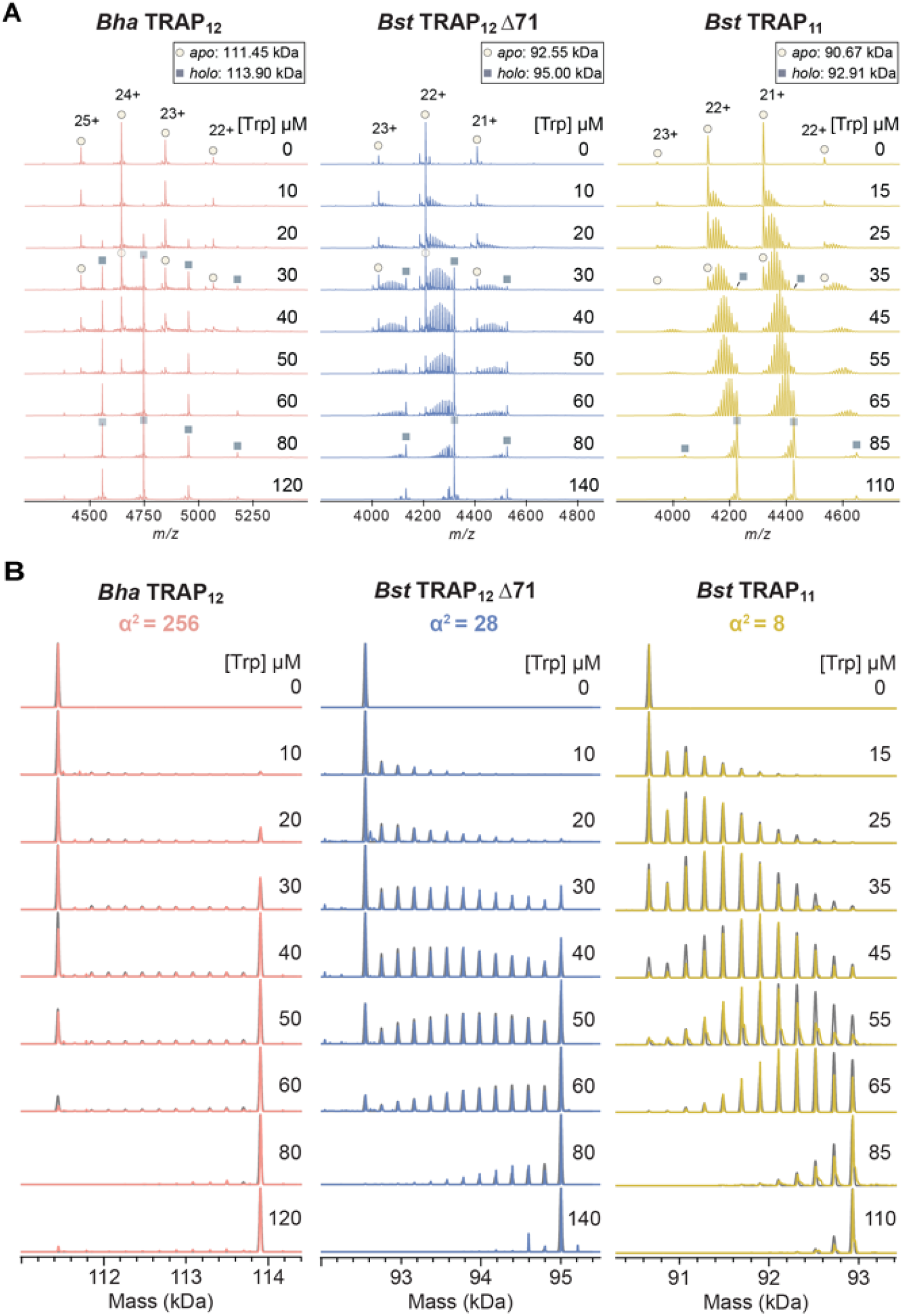
Population distributions measured by native MS reveal extents of homotropic cooperativity in ligand binding to 12mer Bha and Bst Δ71 TRAP (left, middle), and 11mer wild-type Bst TRAP (right). (A) Native mass spectra of the three TRAP oligomers (6.0-7.4 μM Table S3) titrated with 0-140 μM Trp. Increasing Trp in each titration shifts the Trp_n_-TRAP populations from apo (Trp_0_, circles) to holo (Trp_11/12_, squares). Each Trp titration set with either Bha TRAP_12_ (left), Bst TRAP_12_ Δ71 (middle), or Bst TRAP11 (right) features different population distributions, reflecting different degrees of cooperativity. (B) Deconvolved zero-charge mass spectra for the three TRAP oligomers: Bha TRAP_12_ (left), Bst TRAP_12_ Δ71 (middle), or Bst TRAP11 (right). Each experimental dataset (colored) is superimposed with a simulated mass spectrum obtained from best-fit parameters to an additive nearest-neighbor model (grey). Cooperativity is quantified by the parameter α^2^, which is the ratio of the association constant for binding to sites with two bound neighbors, and no bound neighbors (K_N2_/K_0_; figure 2A). Bha TRAP_12_ (left) shows the strongest cooperativity (α^2^ = 256, 1/K_0_ = 718 μM), with populations dominated by apo and holo rings. Cooperativity is moderate (α^2^ = 28, 1/K_0_ = 217 μM) in Bst TRAP_12_ Δ71 (middle) and weak (α^2^ = 8, 1/K0 = 36 μM) in Bst TRAP_11_ (right).

### Differing extent of Trp-Trp cooperativity is evident from population distributions in native MS titrations

Strikingly different population distributions were observed for *Bha*, *Bst* Δ71, and *Bst* TRAP in native MS titrations (Figure 5). For the *Bha* Trp_n_-TRAP_12_ titration, distributions were dominated by either Trp_0_-TRAP_12_ (apo) or fully bound Trp_12_-TRAP_12_ (holo) with minimal population of intermediate Trp_n_-TRAP_12_ states (Figure 5, left). Such a pattern is consistent with strong positive cooperativity (i.e., Figure 2B). Population distributions for *Bst* TRAP were similar to those previously described^16^, with a high population of intermediate states consistent with weak cooperativity. The distributions for *Bst* TRAP Δ71 reflected an intermediate cooperativity between the *Bst* and *Bha* TRAP (Figure 5, middle). In contrast to the highly congruent CD and ITC titrations, the native MS titration data were quite divergent, implying distinctly different underlying thermodynamics.

Fitting of ligand-dependent population distributions revealed strong cooperativity for *Bha* TRAP_12_, moderate cooperativity for *Bst* TRAP_12_ Δ71, and weak cooperativity for *Bst* TRAP_11_. After correcting binding site concentrations using mass-balance (see Methods) the experimental Trp_n_-TRAP_12_ populations were fit with a binding polynomial corresponding to the additive nearest-neighbor (NN-add) model. Datasets were also fit with the non-additive NN model (NN-nonadd), but those resulted in similar thermodynamic parameters and the additional complexity of the model was not justified based on the insignificant improvement in the fits as measured by chi-squared values (see Methods); thus, datasets were analyzed using the simpler NN-add model. For each dataset, we obtained good agreement between populations determined from integration of deconvoluted spectra, and those simulated from best-fit thermodynamic parameters (Figure 5B, Table 1).

**Table 1.** Best-fit thermodynamic parameters for NN-add model of Trp binding cooperativity for native MS titration data (room temperature ~25°C).^†^

| # Bound Neighbors | $\Delta G_{N\#}$ (kcal mol<sup>-1</sup>) | | | $1/K_{N\#}$ (μM)<sup>§</sup> | | | NN cooperativity strength | |
| --- | --- | --- | --- | --- | --- | --- | --- | --- |
| | 0 | 1 | 2 | 0 | 1 | 2 | $\alpha$ | $\alpha^2$ |
| *Bha* TRAP<sub>12</sub> | -4.29 ± 0.12 | -5.93 ± 0.14 | -7.57 ± 0.16 | 720 ± 140 | 45 ± 11 | 2.8 ± 0.8 | 16 ± 5 | 256 ± 87 |
| *Bst* TRAP<sub>12</sub> $\Delta 71$ | -5.00 ± 0.03 | -5.98 ± 0.06 | -6.97 ± 0.09 | 217 ± 13 | 41 ± 5 | 7.8 ± 1.2 | 5.3 ± 0.7 | 28 ± 5 |
| *Bst* TRAP<sub>11</sub> | -6.06 ± 0.20 | -6.68 ± 0.23 | -7.31 ± 0.26 | 36 ± 12 | 13 ± 5 | 4.4 ± 1.9 | 2.85 ± 1.47 | 8.2 ± 4.5 |

<sup>†</sup> Thermodynamic data were obtained by global fitting of titration data to an additive nearest-neighbor thermodynamic model, with parameters for binding to sites with no occupied neighbors $\Delta G$ , and cooperativity terms due to interactions with one or two bound neighbors. In this additive model, the contributions from two neighbors are twice that of one neighbor. Values are best fit from nonlinear least-squares fitting, while standard deviations were determined within three repeats for *Bha*, and *Bst*- $\Delta 71$ TRAP, and two repeats for *Bst* TRAP. Standard deviation was calculated between replicates. <sup>§</sup> Dissociation equilibrium constants are the reciprocal of the corresponding association equilibrium constants.

The best fit parameter values obtained from the additive nearest-neighbor model (Table 1) were consistent with expectations from inspection of the population distribution shifts in the native mass spectra (Figure 5). Cooperativity in *Bst* TRAP yielded a mild eight-fold increase in affinity for sites with two neighbors compared to no neighbors, α^2^ = 8.15, consistent with prior findings.^16^ *Bha* TRAP exhibited remarkably strong cooperativity, with a 256-fold higher affinity for sites with two occupied neighbors. This is consistent with the observation of low abundance for partially loaded states throughout the titration. This is also consistent with the nearest-neighbor model for strong positive cooperativity (Figure 2B). Fits for the engineered 12mer, *Bst* TRAP Δ71 revealed intermediate cooperativity, α^2^ = 28, consistent with the mixed population distributions of partially and fully loaded states (Figure 5). Thus, the nMS experiments, coupled with statistical thermodynamic modeling enabled an accurate description of the thermodynamics of Trp-TRAP binding for variants with cooperativity values differing by a factor of 32 (256/8).

### Microscopic parameters predict macroscopic behavior

Typical methods used for measuring bound fractions (e.g., CD, ITC) yield an average signal that is obtained from all extant micro-states and populations, whereas the NN statistical thermodynamic models are able to define the populations of each individual micro-state. To compare the bulk observable and predicted populations, we simulated macroscopic properties of population distributions of individual protein micro-states over a range of ligand concentrations using the NN-add model and microscopic parameters obtained from fitting the nMS data (Figure 6A, B). These simulations show good agreement between the MS and solution experiments. Discrepancies were largest for the *Bha TRAP* data, whereas they were arguably within experimental error for *Bst* and *Bst*-Δ71 TRAP.

**Figure 6.**
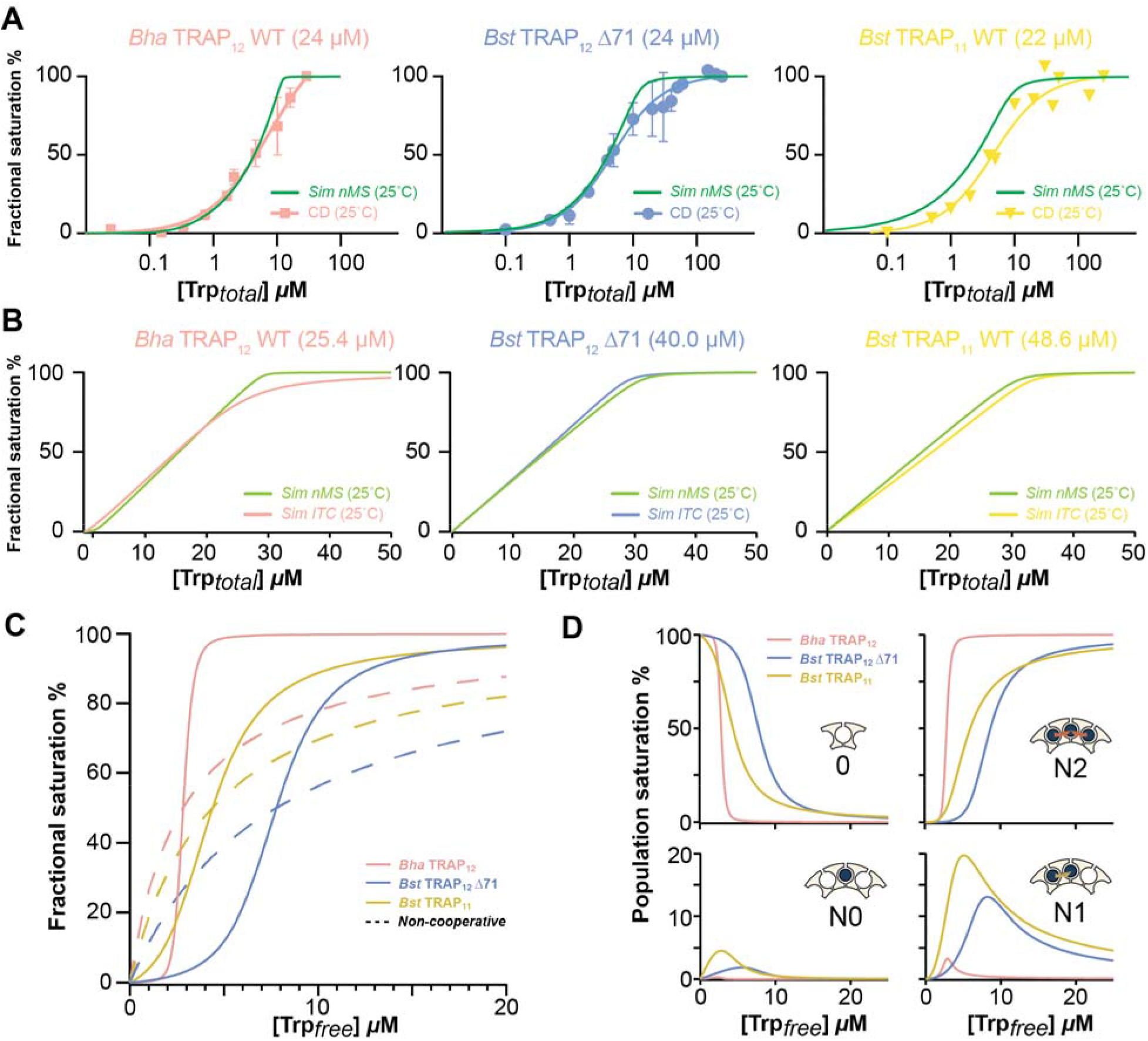
Microscopic parameters from nearest-neighbor cooperativity models can predict macroscopic behavior. (A) Simulated native MS bound fraction from nearest-neighbor parameters (green) and experimental CD signal change (at 228 nm). (B) Simulated native MS bound fraction nearest-neighbor parameters (green) and computed ITC fractional saturation curves based on phenomenological model parameters ΔG and n (Table S2). Simulations used thermodynamic parameters from native MS at room temperature, and temperature and concentrations used in CD and ITC experiments. (C) Simulated mean site occupancy of TRAP proteins over a range of ligand concentrations [Trp_free_]. Bha TRAP_12_ (pink) features a sharp change from almost empty to near complete saturation over the range [Trp_free_] of 2 to 3 μM. Bst TRAP11 (yellow) and Bst TRAP_12_ Δ71 (blue) have more gradual saturation curves due to their weaker cooperativity. Saturation curves with the same apparent K_D_ values in the absence of Trp-Trp cooperativity are shown by dashed lines. (D) Population evolution for the four distinct nearest neighbor binding configurations as a function of [Trp_free_]: (1) empty sites (0), (2) bound sites with no bound neighbors (N0), (3) bound sites with one bound neighbor (N1), and (4) bounds sites with two bound neighbors (N2). The strong cooperativity in Bha TRAP12 (pink) makes N2 the most dominant bound configuration through the range of concentrations. Weaker cooperativity in Bst TRAP_11_ (yellow) and Bst TRAP_12_ Δ71 (blue) allows significant populations of N0, N1, and N2 states over the concentration range. The concentration of populations with adjacent bound ligands can be understood to affect the protein’s regulatory response.

Cooperative behavior in ligand binding is often recognized from sigmoidal isotherms that correlate fractional saturation and free ligand concentration. To approximate this behavior, we simulated fractional saturations over various free Trp concentrations [Trp_free_] (Figure 6C). The cellular concentration of TRAP protein rings in *B. subtilis* cells has been estimated at ~80 nM (0.9 μM Trp binding sites),^36^, while cellular tryptophan concentrations have been measured in the low micromolar range.^37^ Thus, simulations were performed with a binding site concentration of 1 μM and [Trp_free_] to 20 μM. Compared to experimental titrations which monitor response to total ligand concentration (Figure 5), we see the expected sigmoidal curves predicted for positive cooperativity. Despite twenty-fold differences in intrinsic affinities *K*_0_, the apparent affinities fall in a relatively narrow range between ~2 and 8 μM, consistent with those measurements.

We briefly extended the analysis to consider the activation of TRAP for RNA binding. TRAP binds RNA in a Trp-dependent manner, while several binding elements in a single ring are required to achieve high affinity RNA binding.^38,39^ To approximate this behavior, we separately compute bulk saturation and the relative abundance of the four basic nearest-neighbor states (empty sites 0, sites with no bound neighbors N0, sites with 1 bound neighbor N1, and sites with two bound neighbors N2) as a function of free ligand concentration [Trp_free_] (Figure 6D). Consistent with the bulk parameters for *Bha* TRAP (Figure 6C), populations of empty sites drop off steeply in exchange for sites with two bound neighbors, whereas for *Bst* and *Bst* Δ71 TRAP states with one bound neighbor become significantly populated. This behavior can be expected to affect the trajectory of Trp-dependent RNA binding.

## Discussion

### Statistical thermodynamic nearest-neighbor models can readily predict complex ligand behavior in cyclic oligomeric proteins

We used a statistical thermodynamic approach to model the binding of ligands to symmetric ring-shaped homo-oligomeric proteins. By applying nearest-neighbor circular lattice models, we generated partition functions corresponding to the 2^n^ possible liganded states of dodecameric 12mer and undecameric 11mer TRAP rings (Equation 2). Our models considered scenarios in which cooperative interactions between two neighboring ligands are *additive*, meaning simply twice that of having one bound neighbor, and *non-additive*, in which case the effect is not a simple sum of single nearest-neighbor interactions. These parsimonious models allowed us to compute the probability distributions of these states as a function of ligand concentration for a range of cooperativity strengths (Figure 2). Population distributions of states with zero to *n* bound ligands are revealed to be much more sensitive to the underlying microscopic thermodynamic parameters than the bulk metric of fractional saturation. The ability to measure more directly those underlying population distributions can enable us to decipher mechanisms and magnitudes of cooperativity.

### Native MS can directly quantify populations of states with 0 to *n* bound ligands, informing mechanistic binding models

For each of the homo-oligomeric protein variants examined here, native MS enabled us to resolve and quantify populations of oligomers with 0 to *n* bound ligands (Figure 5). These data showed clear and distinct ligand concentration-dependent populations shifts, reminiscent of the population distributions simulated with NN statistical thermodynamic models. Population distributions in each titration series could be fit accurately with an additive nearest-neighbor thermodynamic model *NN-add*, yielding thermodynamic parameters that quantify the affinity of a ligand for isolated sites, *K*_0_, and coupling terms that quantify the allosteric thermodynamic coupling between sites (Table 1). For *Bst* TRAP this coupling energy was 0.62 kcal/mol/neighbor, resulting in an 8-fold higher affinity for binding sites with two neighbors compared to isolated sites, while for *Bha* TRAP a stronger coupling energy of 1.64 kcal/mol/neighbor results in a 256-fold higher affinity. In contrast, conventional ligand binding measurements by spectroscopic and calorimetric methods (Figure 4), failed to clearly distinguish magnitudes of allosteric coupling. Although the population distributions are strikingly different for the three proteins, bulk properties computed from these distributions were in fact similar, suggesting that the native MS measurements have the unique power to resolve underlying population shifts hidden when averaging over all micro-states.

### Implications for understanding of TRAP function

TRAP proteins have evolved to regulate expression of the *trp* operon in response to changing levels of free Trp. Although the TRAP protein sequence is highly conserved,^40^ disparities exist between proteins from organisms that inhabit vastly different environment conditions (i.e., *Bst* – a thermophile, *Bha* – a halophile), in the encoding of the trans-acting protein factor Anti-TRAP (AT),^41^ where the fully transcribed leader possesses sequences capable of preventing ribosome binding to the Shine-Dalgarno sequence^24^, and in their predominant oligomeric states.^42,43^ In the three proteins studied here, two 12mers and an 11mer, we observed orders of magnitude differences in Trp-Trp homotropic cooperativity. The origins of these differences merit additional characterization, but it is tempting to speculate that tighter packing of the otherwise isosteric protomers^22^ results in stronger coupling between sites. Nevertheless, apparent overall affinities of Trp for *Bha* and *Bst* TRAP were similar, possibly reflecting similarities in the fluctuating concentrations of Trp in bacterial cells.

### Limitations

The combination of statistical thermodynamic modeling and native MS to extract and quantify allosteric parameters from the partition function is powerful but has limitations. The most obvious is that while mechanistic models can be readily envisioned for lattices in which nearest-neighbor interactions can be expected to be dominant, such as ring-shaped proteins, it is less straightforward to develop such models for hetero-oligomeric proteins, or homo-oligomers with non-cyclic symmetry elements (e.g., double-rings like GroEL^6^ and the proteosome^44^). Another limitation has to do with the degeneracy of liganded states: e.g., for the NN-add model, the measured populations with half-filled 12mers convolve 924 individual configurations distributed among six distinct statistical weights (*Equation 2*). Other limitations have to do with the measurements themselves: Solution conditions suitable for high resolution native MS typically deviate from the physiological and from solution conditions typical for in vitro biochemical measurements; such deviations could result in context-dependent populations. Moreover, even when native MS-friendly solutions do not perturb populations of states, care must be taken to ensure that ion detection mirrors the populations of states in the solution, avoiding or correcting for differences in ionization efficiencies of different microstates, differences in transmission efficiencies of different ions through the mass spectrometer, or due to potential for dissociation of ligands during ionization and transmission.^18,45,46^ The native MS measurements showed that purified TRAP proteins exhibit additional states, including double-rings and numbers of protomers, and possessed proteofoms (Figure S9, S10). These factors could introduce competing equilibria that complicate efforts to quantify major-state populations in different bound states; of course, such factors would also skew traditional binding measurements that are simultaneously blind to their presence. Nevertheless, with rapidly improving technology for the analysis of proteins by native MS, its combination with statistical thermodynamic models has tremendous potential to advance our understanding of mechanisms of regulation.

## Conclusions

Deciphering mechanisms of allosteric communication between ligand binding sites in oligomeric proteins is difficult. One serious challenge is that for most biophysical measurements the proportionality between the measured signal and populations of liganded states can be distorted by the very property of interest (i.e., allosteric coupling). As an analytical tool, native mass spectrometry has the advantage that the signal does not suffer from this confounding effect – the mass unambiguously tells us the number of bound ligands. As long as the intensity of each signal is proportional to its population, we have an accurate measurement of the underlying distribution of states. Armed with such information, one is then in the position to design and test mechanistic models of site-site communication. In favorable cases, the result can be an understanding of the thermodynamic quantities exchanged between binding sites. These microscopic thermodynamic parameters provide a genuine basis for establishing the structure-thermodynamic relationships responsible for thermodynamic coupling and biological regulation.

## Acknowledgements

This work was supported by National Institutes of Health (NIH) grants R01 GM120923 (to M.P.F. and P.G.), R01 GM077234 (to M.P.F. and P.G.), and P41 GM128577 (to V.H.W.).

We thank Dr. Melody “Pepsi” Holmquist for help with native MS experiments. We also thank members of the Foster lab, Gollnick Lab, and Wysocki lab for useful discussions.

