## Supporting Information for "Thermodynamic coupling between neighboring binding sites in homo-oligomeric ligand sensing proteins from mass resolved ligand dependent population distributions"

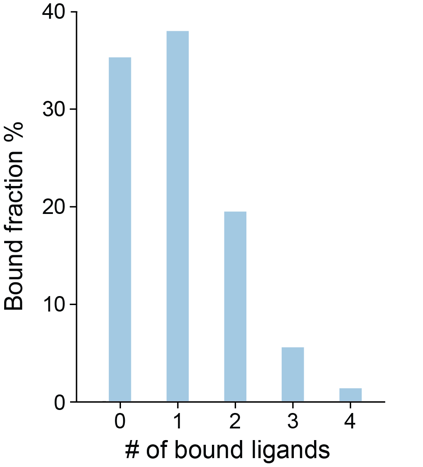

Figure S1. Predicted ligand bound populations of 12mer protein at the saturation level of 1/12 = 8.3% via binomial distribution.

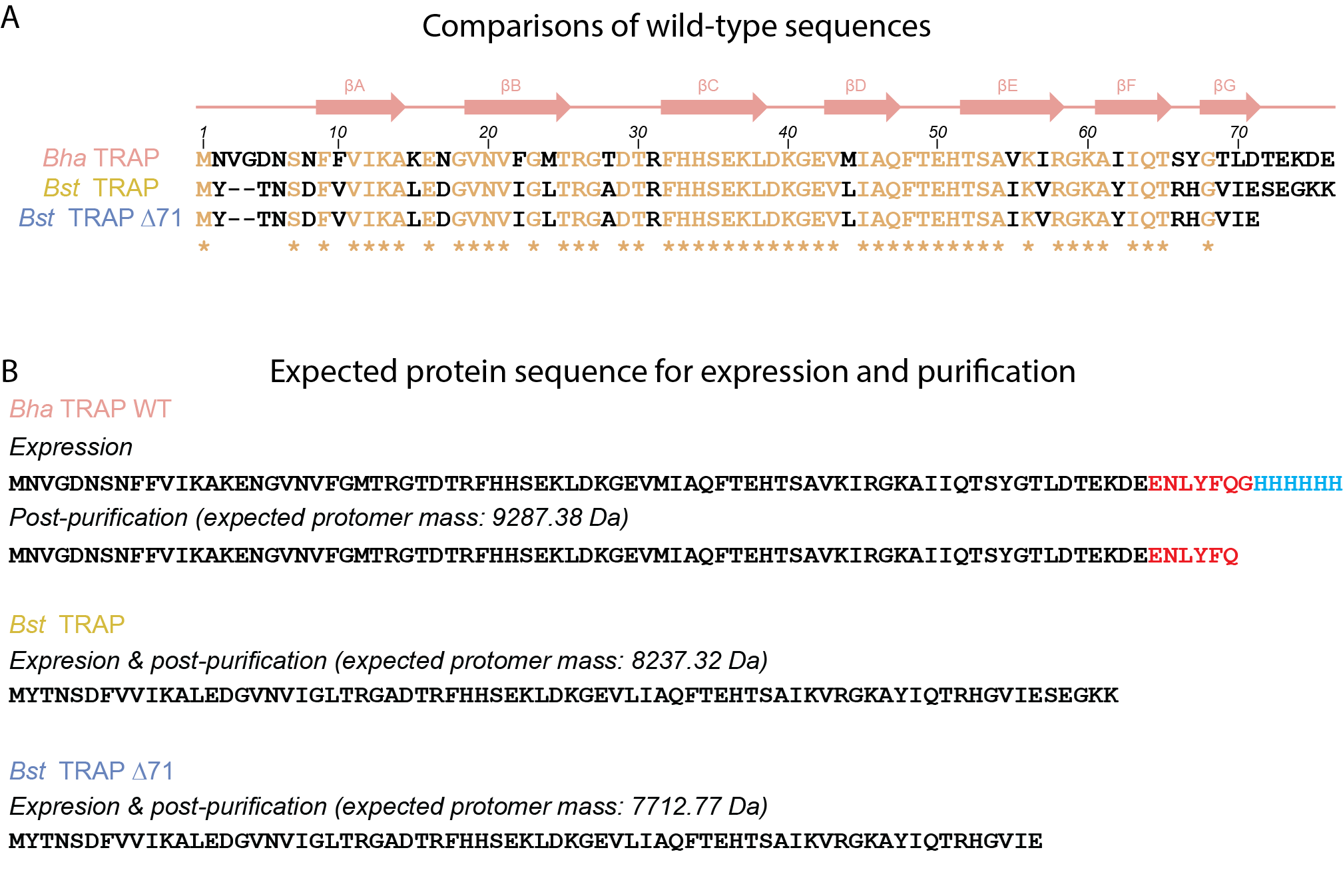

Figure S2. Sequence alignment of TRAP protomers from *Bacillus halodurans* (*Bha*), *Bacillus stearothermophilus* (*Bst*), and *Bst Δ71-TRAP*. (A) The identical residues are labeled with asterisk (*) symbol. Secondary structural elements are from the crystal structure of *Bha* TRAP (PDB 3zzl). (B) Expected protein sequence for expressions and post-purifications with the expected mass.

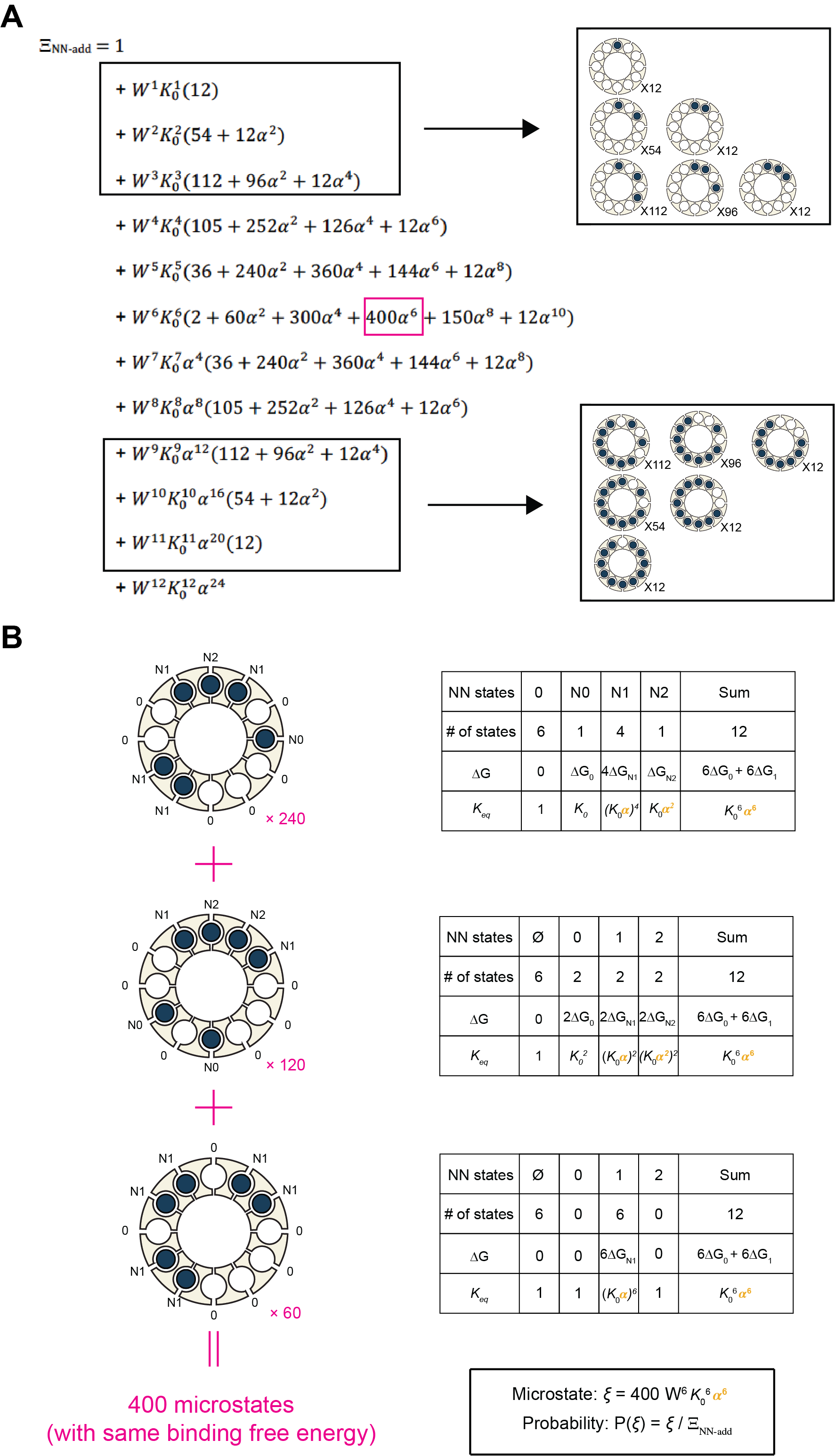

Figure S3. Partition function from additive nearest neighbor (NN-add) model for cyclic homo-12mer protein, and example configurations. (A) Total degeneracies for each macro-state with *m* bound ligands are $W\left( n,m \right)=\left( \begin{aligned} m \\ n \end{aligned} \right)=\frac{m!}{n!\left( m-n \right)}$, where *m* is the total number of sites, and *n* the number of occupied sites. Example configurations are shown for binding with 1, 2, 3, 9, 10 or 11 ligands (rectangles). (B) Detailed nearest-neighbor model illustration of the configurations that give rise to the term 400×W^6^K_0_^6^α^6^ (pink rectangle) in the partition function.

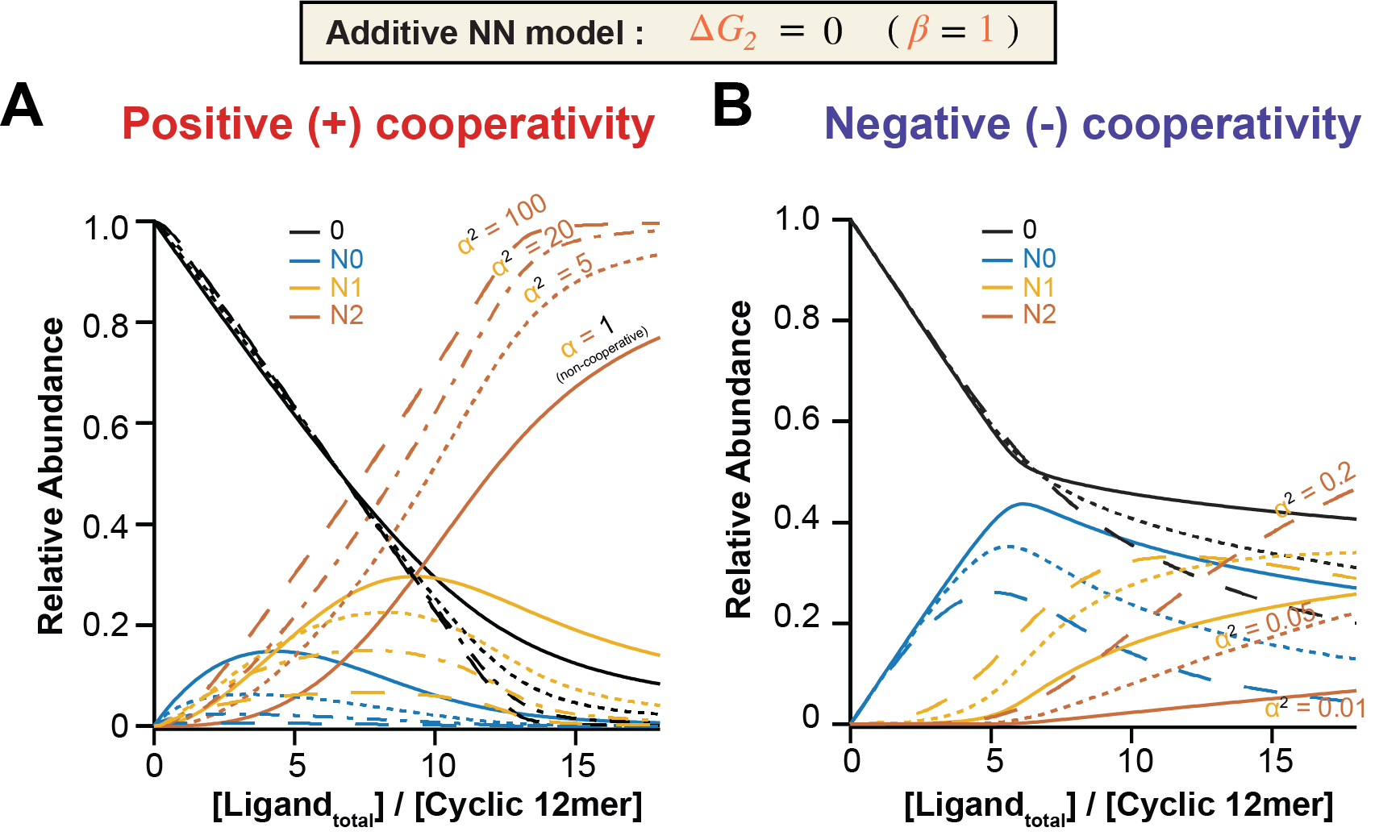

Figure S4. Relative abundance of the four microscopic binding configurations as a function of total ligand concentration as predicted by the additive nearest-neighbor (NN-add) model under different extents of cooperativity (α); simulation parameters were: (A) Cooperativity strengths α^2^ = 1, 5, 20, 100; the non-cooperative case corresponds to α^2^ = 1. (B) Negative cooperativity, with α^2^ = 0.2, 0.05, 0.01. The micro-states are colored differently: empty (black), N0 (blue), N1 (yellow), N2 (brown).

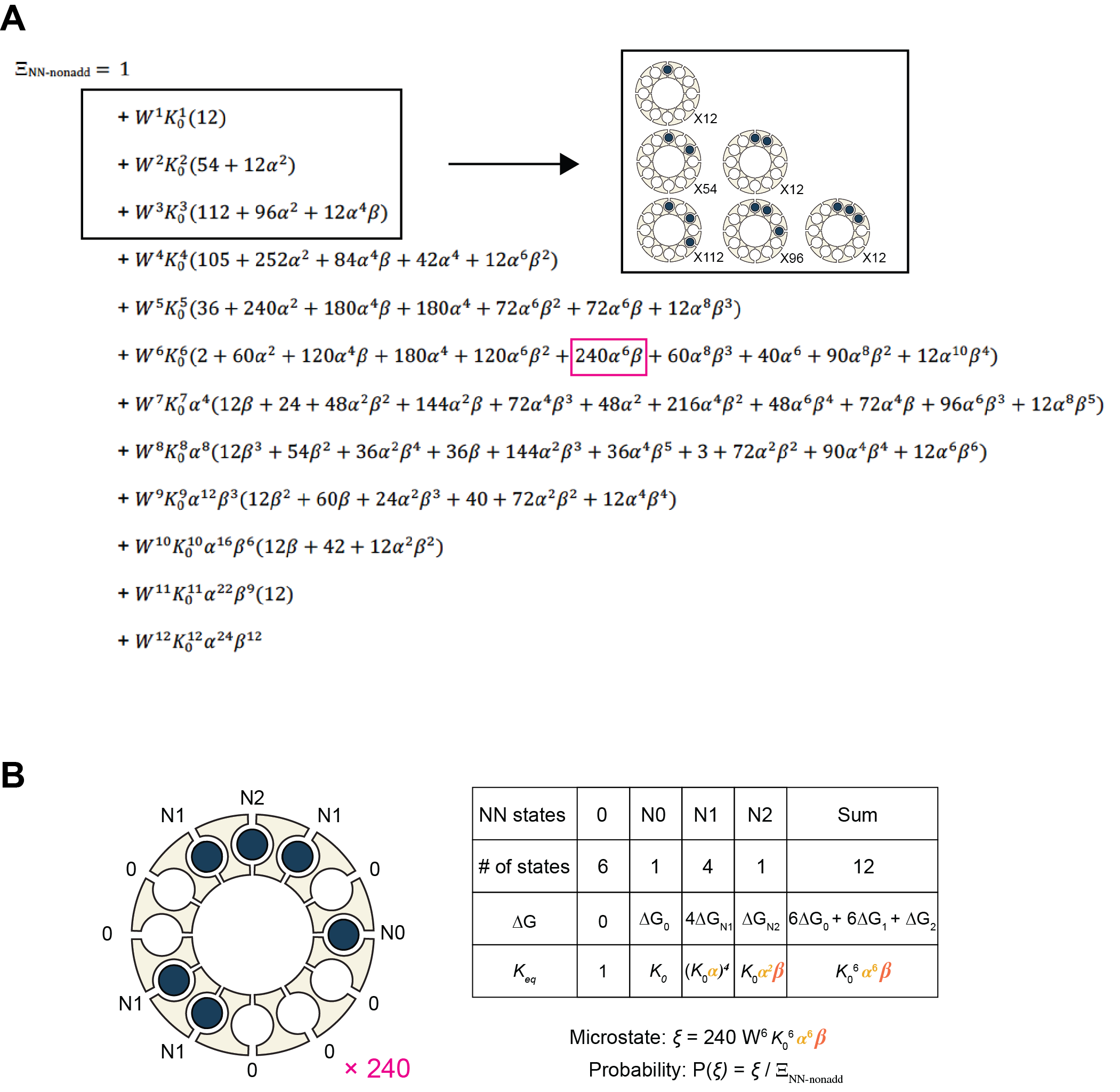

Figure S5. Partition function from non-additive nearest neighbor (NN-nonadd) model for cyclic homo-12mer protein, and example configurations. (A) Example configurations are shown for binding with 1, 2, and 3 ligands (black square). (B) Detailed nearest-neighbor model illustration of the specific configuration of 240×W^6^K_0_^6^α^5^β (pink square) in binding partition function.

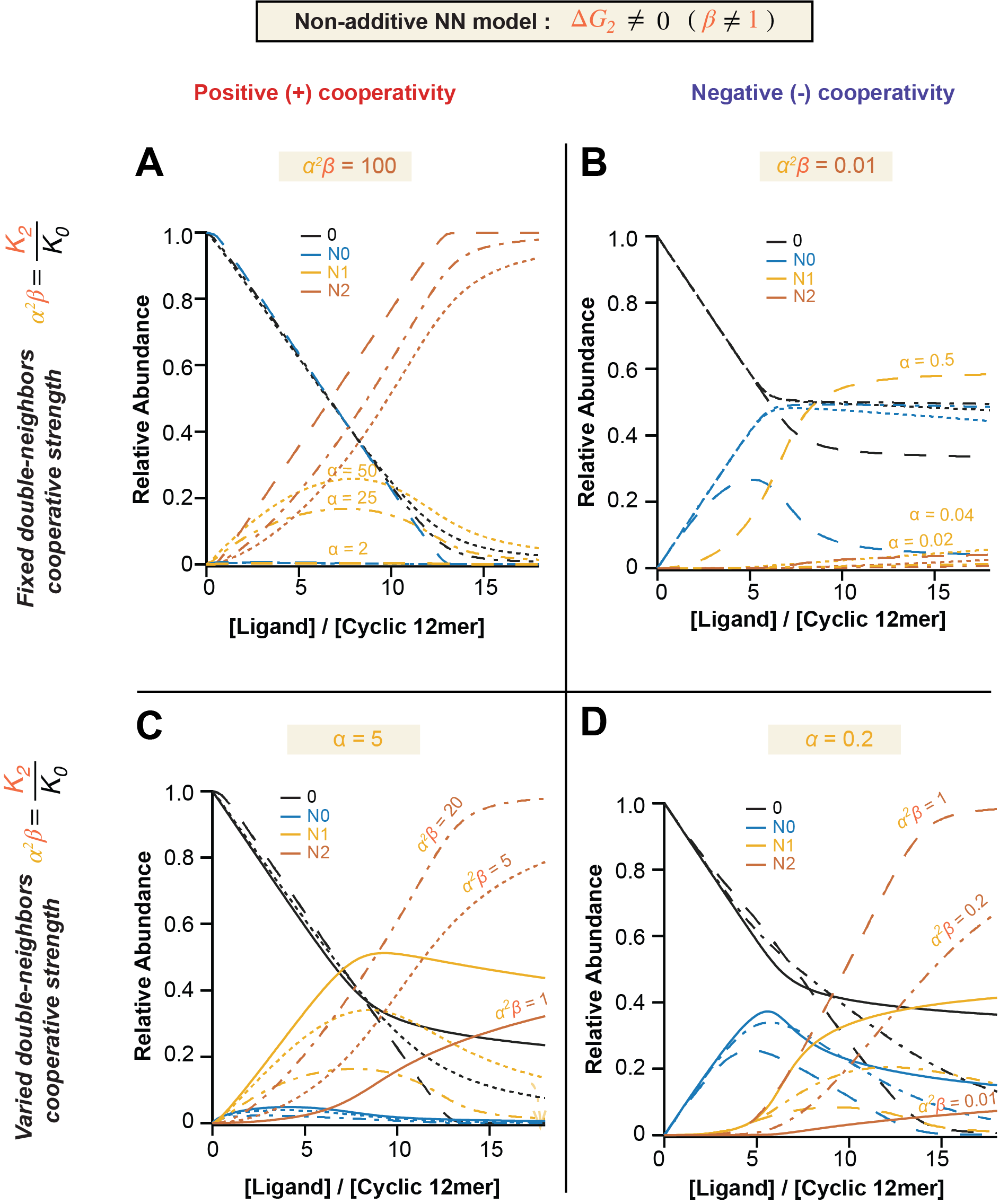

Figure S6. Relative abundance of four basic binding energy configurations in different extends of cooperativity strengths (α and β) predicted by non-additive nearest-neighbor (NN-nonadd) model. Four basic binding energy states are empty (black), N0 (blue), N1 (yellow), and N2 (brown) (Fig. 2A). Plots of relative abundance vs. total ligand / cyclic-12mer ratio were simulated with different α and β, which were corresponding to Fig. 3: (A) positive cooperativity with fixed α^2^β = 100 and varied α and β; (B) negative cooperativity with fixed α^2^β = 0.01 and varied α and β; (C) positive cooperativity with fixed α = 5 and varied β; (D) negative cooperativity with fixed α = 0.2 and varied β.
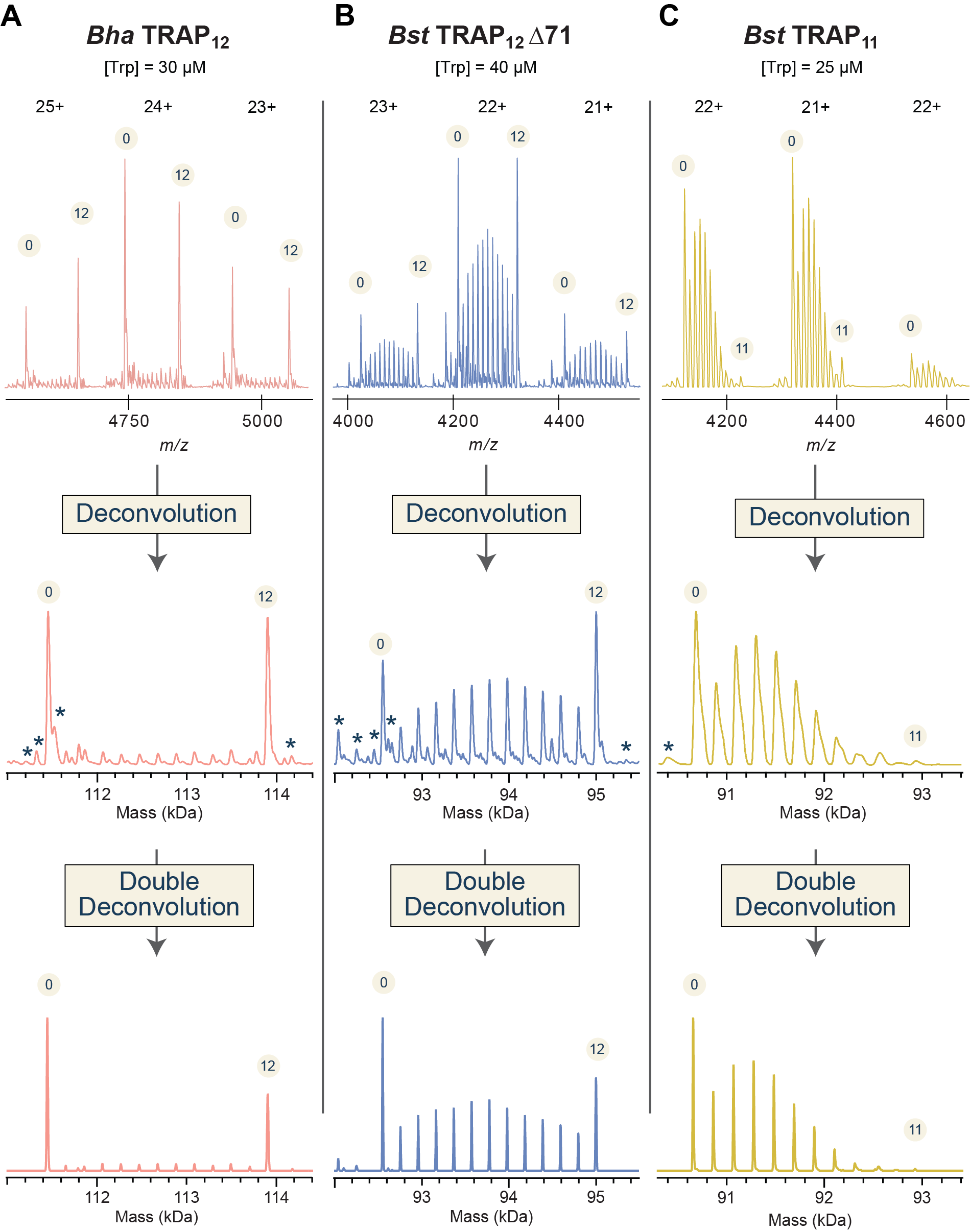

Figure S7. Representative deconvolution processes by Deconvolution and DoubleDec method, sequentially, via UniDec deconvolution program. Only three charge states are shown on the top m/z spectra for each TRAP proteins, followed by the deconvoluted spectra, and double-deconvoluted spectra. Apo and holo states are labeled with circled 0 and 12 (or 11), respectively. Protein proteoforms are labeled with asterisk (*) symbol. (A) *Bha* TRAP mixed with 30 µM Trp. (B) *Bst* TRAP ∆71 mixed with 40 µM Trp, and (C) *Bst* TRAP mixed with 25 µM Trp. Protein concentrations can be found in Table S3.

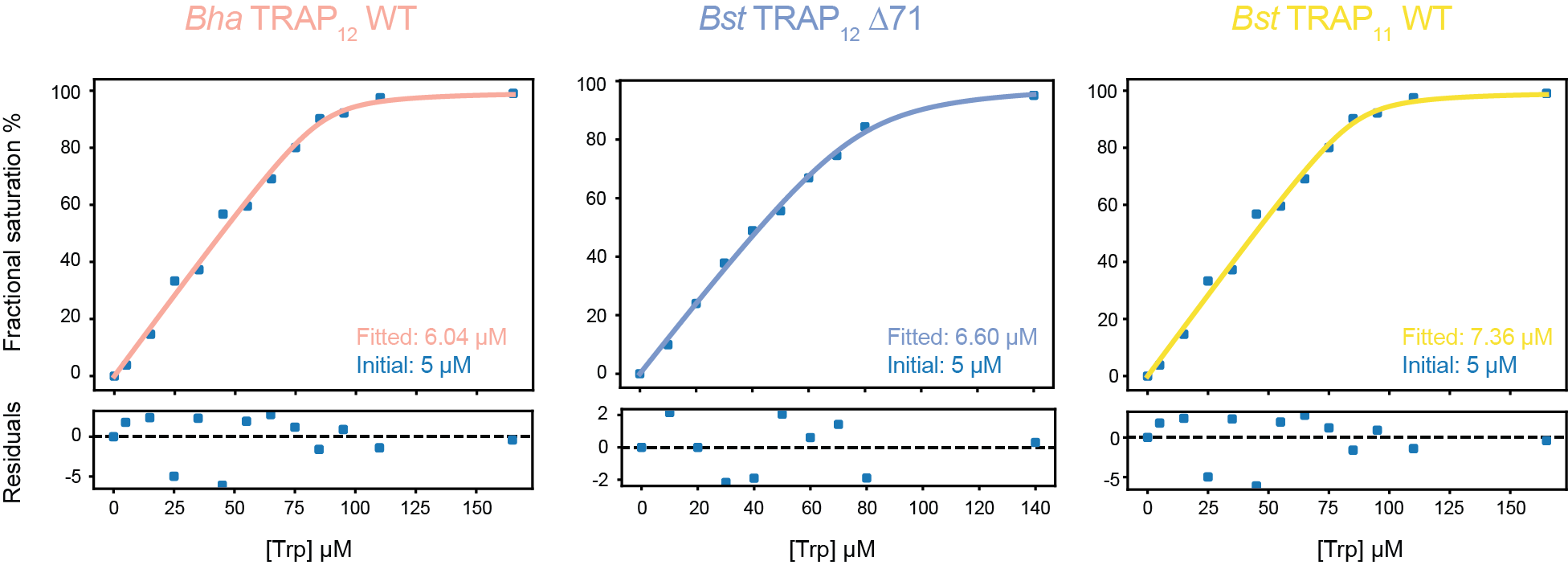

Figure S8. Representative concentration corrections of native MS data using mass balance equation.

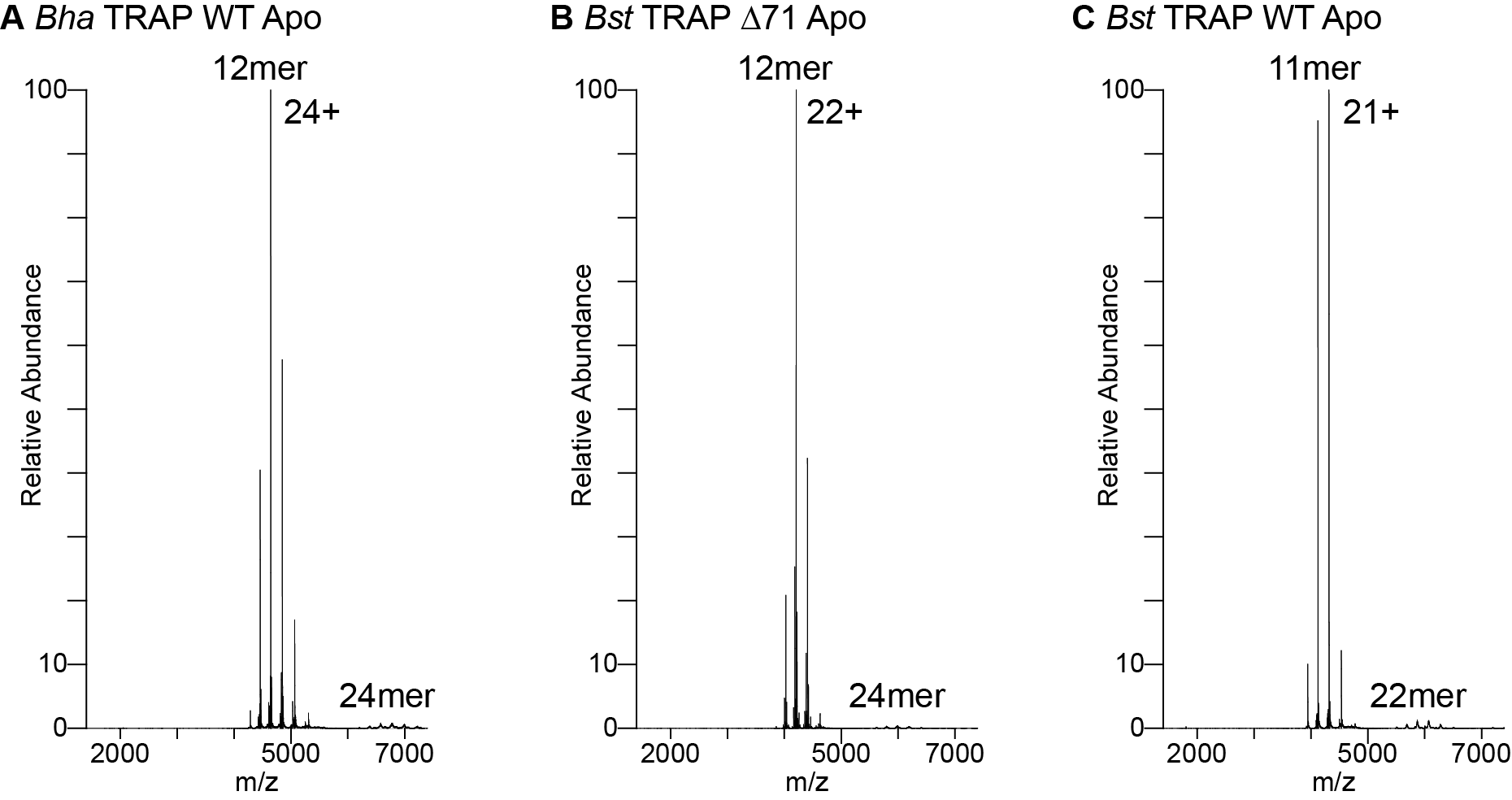

Figure S9. Native mass spectra of the TRAP samples used in this study. *Bha* TRAP (left), *Bst* TRAP ∆71 (middle), and *Bst* TRAP (right) are all at ~ 6 µM ring (Table S3) and in the apo state. The most abundant charge state distribution in each spectrum is the expected oligomeric state for each TRAP sample (labeled the oligomeric state and the most intense charge state). The masses for these oligomers are as follows: *Bha* TRAP_12_ 111444 Da expected and 111453 Da observed, *Bst* TRAP_12_ ∆71 92552 Da expected and 92560 Da observed, *Bst* TRAP_11_ 90662 Da expected and 90670 Da observed. In each spectrum a low relative abundance of “double donut” (24mer or 22mer) is observed. For *Bha* TRAP_12_, there is 13mer overlapping with the expected 12mer charge states.

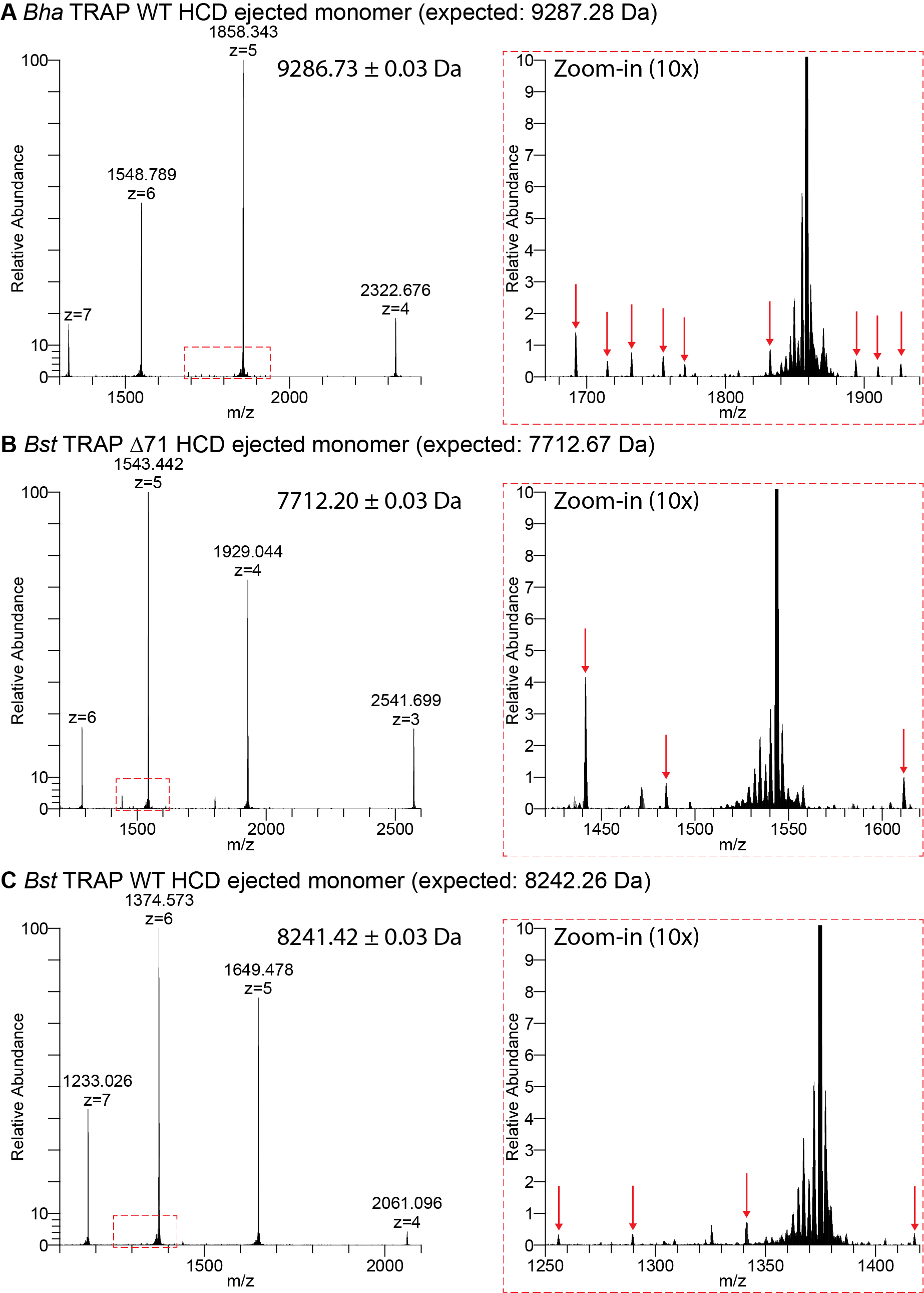

Figure S10. Dissociation (HCD) spectra (left) of (A) *Bha* TRAP, (B) *Bst* TRAP ∆71, and (C) *Bst* TRAP. Zoomed in spectra (right) have red arrows pointing to large mass proteoforms (greater or less than 100 Da).

Table S1 Hill equation analysis of circular dichroism (CD) binding data on Trp-TRAP bindings

|  | ***K_d_* (*µM*)** | **B_max_** |
| --- | --- | --- |
| *Bha* TRAP_12_ | 14.60 ± 1.84 | 128.9 ± 6.4 |
| *Bst* TRAP_12_ ∆71 | 3.97 ± 0.43 | 100.6 ± 1.8 |
| *Bst* TRAP_11_ | 3.02 ± 0.56 | 95.6 ± 3.3 |

Parameters were obtained by fitting CD binding data (37˚C) of Trp-TRAP interactions against the quadratic binding equation (see methods). Values are average ± standard deviation of triplicates for each CD binding data. *K_d_* is the apparent dissociation constant; *B_max_* is the maximum specific ligand binding.

Table S2 Thermodynamic parameters for phenomenological model of Trp cooperativity at 25˚C

|  | **Modes** | | **n** | **∆H**  **(*kcal mol^-1^*)** | **∆G**  **(*kcal mol^-1^*)** | ***K_d_***  **(*µM*)** |
| --- | --- | --- | --- | --- | --- | --- |
| *Bha* TRAP_12_ | Two-sites | Site 1 | 0.70 ± 0.05 | -49.08 ± 3.15 | -7.84 ± 0.10 | 1.79 ± 0.30 |
|  |  | Site 2 | 11.40 ± 0.21 | -8.27 ± 0.17 | -8.31 ± 0.08 | 0.81 ± 0.11 |
| *Bst* TRAP_12_ ∆71 | One-site | | 12.00 ± 0.02 | -18.07 ± 0.07 | -9.60 ± 0.04 | 0.09 ± 0.00 |
| *Bst* TRAP_11_ | One-site | | 11.05 ± 0.03 | -11.47 ± 0.05 | -9.16 ± 0.05 | 0.19 ± 0.02 |

Parameters were obtained by fitting ITC datasets (25˚C) of Trp-TRAP titrations against one-site or two-sites phenomenological binding model. Confidence intervals were obtained by bootstrapping with 200 replicates. *Modes* indicates the type of the phenomenological model. *n* represents fitted binding sites. Dissociation constant from ∆G = RT ln *K_d_* at 25˚C.

Table S3 Corrected [TRAP] and K_d_ via mass balance equation of *DoubleDec*-processed native MS data

|  | **Initial Conc. (µM)** | **Fitted Conc. (µM)** | **Std. Error** | ***K_d_* (µM)** | **Std. Error** |
| --- | --- | --- | --- | --- | --- |
| *Bha* TRAP_12_ | 5 | 6.04 | 0.48 | 0.25 | 0.95 |
| *Bst* TRAP_12_ ∆71 | 5 | 6.60 | 0.56 | 3.12 | 1.98 |
| *Bst* TRAP_11_ | 5 | 7.36 | 0.76 | 0.87 | 3.09 |

Initial concentrations were obtained from protein extinction coefficients at 280 nm. Fitted concentrations were obtained by fitting fractional saturation of Trp binding sites against mass balance equation (see methods) with the assumption of 1-to-1 ratio of Trp binding sites and Trp. Standard errors were estimated using the fits from three replicates of native MS titration data (only two replicates for *Bst* TRAP_11_).
